# Electronic nicotine delivery systems exhibit reduced bronchial epithelial cells toxicity compared to cigarette: The Replica Project

**DOI:** 10.1101/2021.05.05.442767

**Authors:** Massimo Caruso, Rosalia Emma, Alfio Distefano, Sonja Rust, Konstantinos Poulas, Fahad Zadjali, Silvia Boffo, Vladislav Volarevic, Konstantinos Mesiakaris, Mohammed Al Tobi, Antonio Giordano, Aleksandar Arsenijevic, Pietro Zuccarello, Margherita Ferrante, Riccardo Polosa, Giovanni Li Volti, the Replica Project Group

## Abstract

Cigarette smoking is the leading cause of preventable deaths worldwide. Electronic Nicotine Delivery Systems (ENDS) may reduce health risks associated with chronic exposure to smoke and their potential benefits have been the matter of intense scientific debate. Here we replicated three key published studies from the Tobacco Industry on cytotoxic and inflammatory effects of cigarette smoke and ENDS aerosol in an independent multicentric study. We aimed to establish the reliability of results and the robustness of conclusions by replicating the authors’ experimental protocols and further validating them with different techniques. We exposed human bronchial epithelial cell (NCI-H292) to cigarette smoke and to aerosol from ENDS. All the exposure were conducted at air-liquid interface to assess cytotoxicity effects of smoke and aerosol. Moreover, we aimed to assess different inflammatory mediators release (IL-6, IL-8 and MMP-1) from cells exposed to whole smoke and to smoke without particulate matter (vapor phase). We were able to replicate the results obtained in the original studies on cytotoxicity confirming that almost 80% of the cytotoxic effect of smoke is due to the vapor phase of smoke. Moreover, our results substantiated the reduced cytotoxic effects of ENDS aerosol in respect to cigarette smoke. However, our data are significantly different from the original ones in terms of inflammatory and remodeling activity triggered by smoke. Taken all together, the data obtained independently in different laboratories clearly demonstrate the reduced toxicity of ENDS products compared to cigarettes and thus providing a valuable tool to the harm reduction strategies in smokers.

## Introduction

Cigarette smoking is a major risk factor for many pathological conditions, including cardiovascular and respiratory diseases and lung cancer (National Center for Chronic Disease et al., 2014). More than 7 million people die each year using combustible tobacco products, making smoking the leading cause of preventable deaths worldwide (https://www.who.int/news-room/fact-sheets/detail/tobacco). Commercially available innovative combustion-free technologies (e.g., e-cigarettes and tobacco heating products), referred generally as Electronic Nicotine Delivery Systems (ENDS), may reduce the burden of smoking related morbidity and mortality by substantially reducing the health risks associated with chronic exposure to tar from cigarette smoke. The potential benefits and risks of using ENDS have been the matter of intense scientific debate (Bals et al., 2019). A number of toxicological tests are therefore necessary to establish their reduced harm potential compared to combustible cigarettes and to ensure protection of individual and public health from the adverse effects of potentially harmful exposures (Krewski et al., 2020).

In 2007, the Committee on Toxicity Testing and Assessment of Environmental Agents (Krewski et al., 2020; Russell & Burch, 1959) proposed a strategy for toxicity testing. In particular, the goal of toxicity testing has been established as aimed to identify pathways that, when perturbed, can lead to adverse health outcomes and to evaluate the host susceptibility to understand the effects of such perturbations on humans. Among the most important pre-clinical studies on the effect of ENDS with respect to tobacco cigarette smoke is the studies conducted by the R&D of Tobacco Companies. Stand out studies evaluated the cytotoxicity induced by smoke and aerosol on cultured cell models of human bronchial epithelium (Azzopardi et al., 2016; Gonzalez-Suarez et al., 2016; Iskandar et al., 2017), endothelial cells (Anderson et al., 2016; Taylor et al., 2017), immune cells (Poussin et al., 2015), as well as studies on inflammation (Azzopardi et al., 2015; Xu et al., 2019), oxidative stress (Rayner et al., 2021; Taylor et al., 2016), genotoxicity and mutagenicity (Rudd et al., 2020).

Herein we focused on key published studies from the Tobacco Industry, exhibiting a significant impact on policies and public health. In particular, we aimed to replicate the findings of such key studies by carefully replicating the methodological approach used by the authors and by integrating the experimental protocol with alternative scientific methods.

This replication study is mainly based on the reproduction of the original study results, but also on the reliability of measurements, verifiability of results and robustness of conclusions. In order to improve the reproducibility of our results and to minimize bias, we planned a multi-site study (ring study).

The main goal of this study was to replicate and confirm three “high impact” published works about cytotoxicity induced by ENDS aerosols and tobacco smoke on a model of airway epithelial cells (NCI-H292) (Azzopardi et al., 2015; Azzopardi et al., 2016; Jaunky et al., 2018). These papers assessed the cytotoxic effect induced by the smoke from a reference cigarette, 3R4F (University of Kentucky), the aerosol from two electronic cigarette (Vype e-Pen and Vype e-Stick) and the aerosol from two Tobacco Heating Products (IQOS and GLO™) in the same human bronchial epithelial cell model (NCI-H292) and with the same exposure system at air-liquid interface (ALI). In particular the first paper (Azzopardi et al., 2015) assessed the cytotoxicity induced by both whole smoke (WS) and vapor phase (VP), produced by 3R4F cigarettes with two different regimes: International Organisation for standardization (ISO) and Health Canada Intense (HCI). The VP was represented by the whole smoke deprived of the total particulate matter (TPM) placing a Cambridge Filter Pad (CFP) in line between the reference cigarette and the exposure chamber. Moreover, this work evaluated the different ability of WS and VP to induce inflammation and remodeling by measuring the production and release of IL-6, IL-8 and MMP-1 in the media at 24 hours from the exposure to smoke. The second paper by Azzopardi et al. (Azzopardi et al., 2016) aimed to assess the cytotoxic effects of two e-cig (Vype e-Pen and Vype e-Stick), and the third paper by Jaunky et. al (Jaunky et al., 2018) aimed to evaluate the cytotoxic effect of two THPs (IQOS and GLO™) in the same cell model (NCI-H292) and by the same ALI exposure system. We decided to replicate the most relevant findings of these three studies about cytotoxicity, expanding the findings on inflammatory cytokines induced by WS and VP from tobacco cigarette of the first study (Azzopardi et al., 2015) to the comparison between the two different smoke regimes (ISO and HCI).

## MATERIALS AND METHODS

### Recruitment of Laboratories

International Laboratories involved in cytotoxicity studies were selected to participate in the inter-laboratory “Replica” study based on predefined acceptance criteria. Before the start of the study, we provided access to an online questionnaire which listed a body of skills and knowledges pertaining to the core activities of *in vitro* research in order to assess levels of proficiency in general and to specific areas of our research, including experience in biological assessments of cytotoxicity and ELISA determination. A section of the questionnaire outlined required equipment and “smoke” laboratory compliance according to ISO3308:2012 (International Organization for Standardization, 2018), GLP and EPA guidelines. Workshops, hands-on training and on-site assessment of laboratory capacity and personnel expertise were established thereafter and follow up virtual sessions were conducted when necessary. In particular dedicated scientists with varying levels of experience in *in vitro* testing were trained in the critical phases of exposure. The majority of them did not have previous formal training in smoke and aerosol exposure procedures. They were subsequently trained in the proposed SOPs for smoking/vaping machines utilization and smoke/aerosol cell exposure. In total, 4 laboratories from the academia and one from the private sector were selected and joined this study, respectively from: Italy (LAB-A), Greece (LAB-B), Oman (LAB-C), USA (LAB-D) and Serbia (LAB-E).

### Standardized Operative Procedures (SOP) Development and Harmonization Process

We strengthen the significance of our collaboration by harmonization of laboratory protocols, defining Standard Operating Procedures (SOPs) for each experimental step, using the same cell line, the same cell-exposure systems (e.g., smoking machine and vaping machine) and the same methods to assess endpoints. Many of these actions are suggested by the Transparency and Openness Promotion (TOP) guidelines (https://www.cos.io/initiatives/top-guidelines). A four-day kick-off meeting was held by the leading center in Catania, Italy (Lab A). Each meeting was split in two sessions. One for harmonization of Standard operating procedures (SOPs) and the other for personnel training. SOPs for cell exposure to cigarette smoke and ENDS aerosol, cell culture, cytotoxicity assessment using the neutral red uptake (NRU) assay and ELISA cytokines determination were adopted from replicated papers and manufacturer instructions. The original version of SOPs was revised by the leading center, then particularized with the principal investigators site and adapted to laboratory capacity, equipment and test products and following ISO3308:2012 guidelines (International Organization for Standardization, 2018). A scientist from each laboratory was assigned to participate into a theoretical-practical training course in the laboratories of the leading center (LAB-A). The training focused on the functioning of smoking/vaping machines, exposure of the cells to smoke/aerosol, execution and reading of the NRU assay. Datasheets for detailed recording of technical data related to critical protocol steps and deviation communication forms were prepared and shared with laboratory partners, in order to provide corrective actions in case of deviations in laboratory results. For data analysis and reporting, spreadsheet (Microsoft Excel) templates were prepared and distributed to the laboratories. These documents were collected in individual folders with restricted access to one partner only and the data manager. After this phase and at completion of equipment set-up at each center, both the Principal Investigator (PI) and co-Principal Investigator (co-PI) planned a visit each to the participating laboratories to verify the correct installation and functioning of the smoking Labs, to assess compliance to standards and to conduct additional on-site training in the partner laboratories. Unfortunately, due to the restrictions imposed by the SARS-CoV-2 pandemic, it was not possible to carry out on-site training in the US laboratory (Lab D), so in this case three different remote visits were conducted via video conferences. To ensure better assay standardization of cell growth, cytotoxicity assessment and cytokines determination, a list of key consumables was outlined and shared with all laboratories. In particular, the same lot of fetal bovine serum and test products were provided to each laboratory. Human NCI-H292 bronchial epithelial cell line was purchased (ATCC, CRL-1848). A harmonized SOP for thawing, freezing and subculturing of the NCI-H292 cell line, and testing for mycoplasma contamination (Plasmotest™; InvivoGen) as an essential quality control before freezing the cells, was used by the study partners to generate their own working cell bank.

### Study design

The study was designed in three stages. In the first stage we assessed the IC50 for NCI-H292 cells exposed to smoke (WS and VP) using two different regimes (ISO and HCI), as per Azzopardi et al. (Azzopardi et al., 2015). In the original study, the authors used a Borgwaldt RM20S smoking machine (Hamburg, Germany) equipped with 20 channels with two different syringes per channel, able to dilute smoke with air at each puff with a predetermined dilution factor ranging from 1:500 to 1:2.5 (smoke:air, vol:vol) thus creating a cell viability curve based on the dilution factor of cigarette smoke with air. In our study we used a Borgwaldt LM1 smoking machine able to convey only undiluted smoke on cells. For this reason, while Azzopardi et al in their work calculated the EC50 for viability of NCI-H292 cells, we calculated the IC50, which is more convenient for undiluted smoke exposure. At this stage we had to define our cell viability curve based on puff number and nicotine released in the culture media, in order to identify the IC50 of cells by our exposure system, and to compare the effects between WS *vs*. VP produced by 1R6F cigarette. During the second stage, we replicated another study by Azzopardi et al. (Azzopardi et al., 2016) reproducing the same experiments from the first stage, but exposing cells to electronic cigarettes aerosol and comparing the cytotoxic effects to 1R6F cigarette whole smoke. In the third stage we replicated a study by Jaunky et al. (Jaunky et al., 2018) by exposing NCI-H292 cells to aerosol from two THPs, using the same previous conditions, and comparing their cytotoxic effects with 1R6F cigarette WS and e-cigarettes aerosol. For each exposure, basal media was collected from the bottom of the exposure chambers, and nicotine dosimetry was conducted by Lab A using UPLC-ESI-TQD analysis. Exposure with both e-cigarettes and THPs was conducted at a number of puffs based on their respective IC50 values of 1R6F WS obtained using the HCI regime exposure, in order to perform comparable cell exposures with the different products. Finally, at each stage, measurements of concentration of secreted inflammatory (IL-6 and IL-8) and tissue remodeling (MMP-1) mediators were assessed for 1R6F dose response curves (WS *vs*. VP).

### Test products

In the original studies 3R4F cigarettes (University of Kentucky) were used to expose NCI-H292 cells to cigarette smoke. 3R4F cigarettes are no longer produced and have been replaced with 1R6F cigarettes (University of Kentucky) (Jaccard et al., 2019), which we adopted as reference for our exposures. As per original papers, 1R6F cigarettes were conditioned for a minimum of 48 h prior to use (60 ± 3% relative humidity, 22 ± 1 °C) and smoked in a test atmosphere of 60 ± 5% relative humidity, 22 ± 2 °C in accordance with ISO 3402:1999 (International Organization for Standardization, 1999). Cigarettes were either smoked according to ISO 3308:2000 (35 mL puff volume, drawn over 2 s, once every minute with ventilation holes unblocked) or to the HCI smoking regimen (55mL puff volume, drawn over 2s, twice a minute with ventilation holes blocked). Moreover, two commercially available e-cigarettes, the Vype eStick and a Vype ePen 3 (Nicoventures, Blackburn, UK; www.govype.com) were used in this study. In the original paper by Azzopardi et al (2016) was used the first version of the Vype ePen, which is no longer available and has been replaced with Vype ePen3. Vype ePen3 is a button-activated “closed-modular” system consisting of two modules, a rechargeable battery section 650 mAh power with 6-Watt resistance (output of 5.0V) and a replaceable liquid (“e-liquid”) containing cartridge (“cartomizer”) equipped with a cotton wick heating system. Vype eStick is a puff-activated cigarette-like product consisting of two modules, a battery unit with a capacity of 280 mAh and a pre-filled cartridge containing e-liquid and cartomizer. We used the “Master Blend” flavor for Vype ePen3 and the “Toasted Tobacco” flavor for Vype eStick, both with 18 mg/ml nicotine. Finally, two commercially available THPs, GLO™ (named as THP1.0 in the paper by Jaunky et al.) and IQOS (named as THS in the paper by Jaunky et al.), were tested at the same puffing regime (HCI). Jaunky and colleagues used the products named GLO™ and IQOS, instead we used the newest generation at the time of this study: GLO™ PRO (https://www.discoverglo.com) and IQOS 3 DUO (https://iqos.com/). For GLO™ PRO we used the Neostick “Ultramarine”, and for IQOS 3 DUO we used the “Sienna selection” (Red) Heets. All devices and consumables for test products were sourced from Italy, except for the Vype eStick, which was kindly provided by Greece.

### Smoke and aerosol generation and exposure parameters

The Borgwaldt LM1 Smoking Machine and the LM4E Vaping Machine (Borgwaldt, Hamburg, Germany) are able to expose cells respectively to undiluted smoke and aerosol. Cell culture exposure chambers used in this study were previously described by Azzopardi and Jaunky (Azzopardi et al., 2015; Jaunky et al., 2018). In the first phase we exposed cells to 1R6F smoke using the two regimes, the ISO 3308:2000 (puff volume, duration and frequency of 35 mL, 2 s and 60 s (35/2/60), with bell shape) (International Organization for Standardization, 2000) and the Health Canada Intense (puff volume, duration and frequency of 55 mL, 2 s and 30 s (55/2/30), with bell shape) smoking regimes. These exposures were conducted both to whole smoke (WS) and to vapor phase (VP) of smoke, obtained positioning a Cambridge Filter Pad (CFP) in line, immediately after the cigarette. Based on nicotine concentration released at IC50 dose of HCI WS exposure, we have set the puff number able to reach the same nicotine concentration in the exposure chamber media in order to perform the exposure of NCI-H292 cells to e-cigs and THPs aerosol. Vype ePen 3 was vaped for 10 puffs using a modified HCI regime (mHCI; puff volume, duration and frequency of 55 ml, 2 s, 30 s, (55/2/30) and with a rectangular shape) with 1 second of pre-activation for each puff (Azzopardi et al., 2016). Vype eStick was vaped for 25 puffs using the CORESTA Reference method n. 81 (CRM81) regime (puff volume, duration and frequency of 55 ml, 3 s, 30 s, (55/3/30) with rectangular shape). THPs were manually button-activated to initiate device heating prior to syringe activation. GLO™ PRO was activated 30 s prior to puffing and the Neostick was puffed 8 times; IQOS 3 DUO was activated 20 s prior to puffing and the Heets was puffed 7 times. Different heat cycles were mandated by the product design specification for each THP, as described in the manufacturers’ usage instructions. THPs were vaped following the HCI regime (puff volume, duration and frequency of 55 mL, 2 s and 30 s (55/2/30), with bell shape), but with filter vents unblocked to avoid device overheating. Prior to analysis, 1R6F cigarettes were conditioned for at least 48h at 22±1_C and 60±3% relative humidity in accordance with ISO 3402 (ISO, 1999). E-cigarettes were fully charged and loaded with fresh cartomizers for each exposure. Vype ePen and eStick were held at a 45° angle (mouthpiece up), reflecting observed consumer use of the product. THPs were fully charged, cleaned and loaded with fresh Heets or Neosticks for each exposure.

### Chemicals and reagents

Chemicals and reagents were obtained from the following sources: Dulbecco’s Modified Eagle Medium (D-MEM), RPMI-1640 medium (w/o glutamine), phosphate buffered saline (PBS), Penicillin–Streptomycin solution 10.000 U/ml, L-glutamine 200 mM, Transwell® culture inserts (12 mm diameter, 0.4 μM pore size) and trypsin–EDTA from ThermoFisher Scientific; glacial acetic acid, neutral red solution, formaldehyde solution and Absolute ethanol (≥99.8%) from Sigma–Aldrich™; fetal bovine serum - Sud America Origin (FBS: Corning; LOT#35079016) and UltraCULTURE™ from Lonza (Basel, Switzerland); interleukin-6 and -8 (IL-6 and IL-8) human Instant Elisa™ kits and matrix metalloproteinase-1 (MMP-1) human Elisa kit from ThermoFisher Scientific.

### Cell culture

The NCI-H292 human bronchial epithelial cells from the American Type Culture Collection (ATCC; cell no. CRL-1848) were chosen to replicate the original studies (Azzopardi et al., 2015; Azzopardi et al., 2016; Jaunky et al., 2018). These cells were chosen as a cell model representing the respiratory tract directly exposed and a major site of deposition of smoke and vapor respectively from tobacco cigarettes and ENDS. Moreover, these cells are easy to maintain, and therefore suitable for standardization among the different laboratories. Briefly, NCI-H292 cells were cultured in RPMI 1640 medium (10% FBS, 2 mM L-glutamine, 50 U/ml penicillin and 50 mg/ml streptomycin) at 37 °C, 5% CO_2_ in a humidified atmosphere. Then, cells were seeded in 12 mm Transwell® inserts (Corning Incorporated, NY, USA) at a density of 3×10^5^ cells/ml sustained by 1 ml of RPMI medium in the basal compartment of each well and 0.5 ml in the apical compartment of each Transwell® insert, 48 hours prior to exposure. Cell starvation was done 24 hours prior to exposure by replacing the basal and apical medium with 1 mL and 0.5 mL respectively of UltraCULTURE™ containing

2 mM glutamine, 50 U/mL penicillin and 50 μg/mL streptomycin. When the 80% confluency was reached, the apical medium was removed from each insert and two inserts per test product were transitioned to the exposure chamber with 20 ml of DMEM-high glucose (DMEM-hg) in the basal compartment in order to perform the air-liquid interface (ALI) exposure (**Fig. 1**). For each smoking/vaping regime, one exposure chamber was connected to a LM4E port without the ENDS device so as to expose cells to laboratory air filtered by a Cambridge Filter Pad at the same regime (AIR control). Moreover, 2 negative controls, consisting of 1 seeded insert with apical media (INC) and 1 seeded insert without apical media (ALI) in the incubator, and 1 positive control with 1 ml apical and 2 ml basal sodium dodecyl sulphate (SDS) at 350 μM were included for each exposure run. After each exposure, the inserts were transferred from the chamber to a clean well plate, adding 1 mL and 0.5 mL of supplemented UltraCULTURE™ respectively at the basal and apical side for 24 hours of recovery period.

**Fig. 1.**
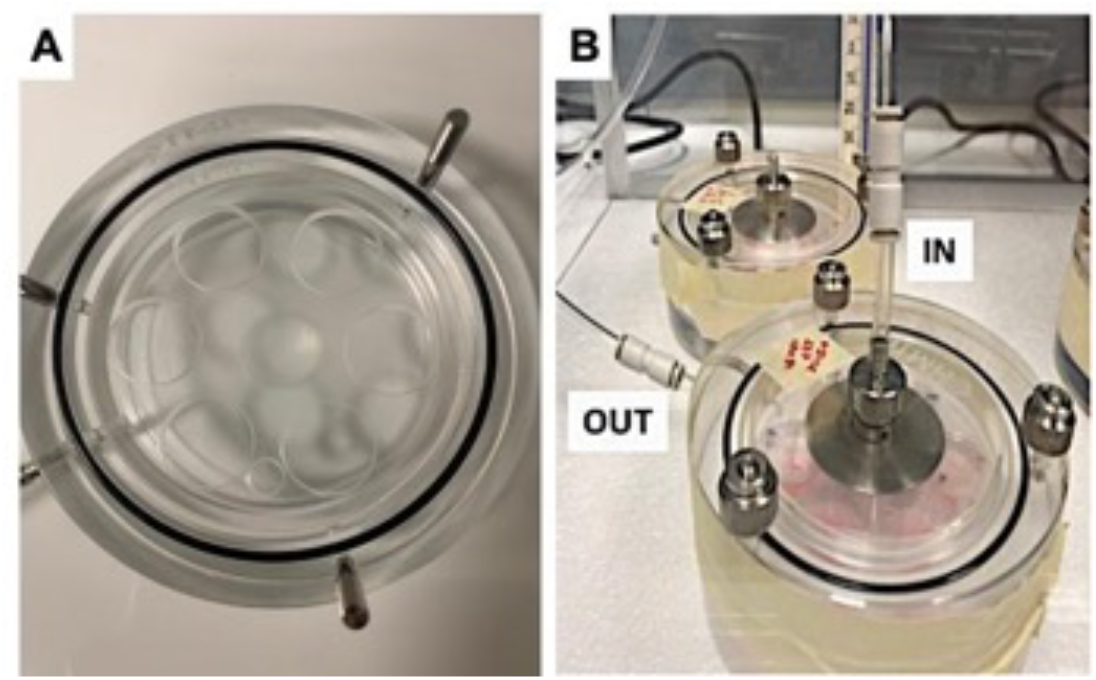
Perspex aerosol exposure chamber

### Nicotine dosimetry

Nicotine dosimetry was performed on 0.1 ml aliquots of media contained in the exposure chambers. Each sample and calibration standard (1, 2, 5, 10, 20, 50 µg/ml) were added with nicotine-(methyl-d3) solution used as internal standard at 100 µg/ml. Subsequently, 0.1 ml of sulfuric acid 0.1 M and 0.3 ml of acetonitrile were added to each sample, all of which have been vortexed and centrifuged at 2500 RCF for 4 min. Afterwards, supernatants were transferred to vials with a 250 μl conical insert. Nicotine was determined by UPLC-ESI-TQD (Waters Acquity), operating in Multiple Reaction Monitoring (MRM) and positive ion mode. An Acquity UPLC® HSS T3 1.8 μm – 2.1×100mm column was used. Isocratic elution (80% water and 20% acetonitrile, both added at 0.1% with formic acid) was performed. The mass spectrometry settings were as follows: capillary energy at 3.0 kV, source temperature at 150 °C, column temperature at 40 °C, desolvation temperature at 500 °C, desolvation gas at 1000 L/hr and cone gas at 100 L/hr.

**Table 1.** MRM transitions monitored (m/z) with cone and collision voltages.

| **Analyte** | **MRM (*m/z*)** | **Cone (volts)** | **Collision energy (eV)** |
| --- | --- | --- | --- |
| Nicotine | 163 → 117 | 40 | 25 |
|  | 163 → 132 | 40 | 15 |
| Nicotine-(methyl-d3) | 165.8 → 116.8 | 40 | 20 |
|  | 165.8 → 129.7 | 40 | 20 |

### Tobacco whole smoke (WS) and vapour phase (VP), ENDS aerosol and air exposure of cells

The smoke exposure system used in this study has been previously described (Adamson et al., 2011; Azzopardi et al., 2015; Jaunky et al., 2018; Maunders et al., 2007; Phillips et al., 2005; Thorne et al., 2009). Cells, prepared 24 h prior to exposure, were transitioned to the ALI by removal of the covering apical cell culture medium, transferred into a perspex aerosol exposure chamber (Fig. 1) and then exposed to different puffs by ISO regime (2, 5, 10, 12, 15, 20, 25, 30 puffs) and by HCI regime (2, 4, 5, 6, 8, 10, 15, 20 puffs) of mainstream WS or VP using a Borgwaldt LM1 smoking machine (Hamburg, Germany; **Fig. 2**). VP exposure was achieved by the addition of an in-line Cambridge filter pad (CFP) to remove total particulate matter (TPM). During exposure, cells were fed basally with DMEM containing 50 U/mL penicillin and 50 μg/mL streptomycin. Throughout WS and VP exposure, cell cultures were maintained at 37 °C in a thermostatic incubator close to the smoking machine. Following exposure, the culture inserts were transferred back to fresh 12-well culture plates containing 1 mL supplemented UltraCULTURE™ pre-warmed at 37 °C. 0.5 mL of supplemented UltraCULTURE™ was added to the apical surface of each culture insert and the cells incubated for a further 24 h at 37 °C, 5% CO2 in a humidified atmosphere. Control cultures, in which culture medium covering the apical surface of the cells was removed, were either returned to the incubator (ALI) for the same time of the exposure of samples, or exposed to a flow of filtered (by CFP) laboratory air at the same regime and max puff number of the samples (AIR). An incubator control (INC) was included in which cells were maintained submerged in culture medium at 37 °C and 5% CO_2_ throughout the exposure and 24 h recovery period. Following the recovery period, culture medium from the apical and basal compartments of each culture inserts and from all exposure studies were individually pooled and stored at −80 °C until the secreted inflammatory and tissue remodeling mediators were measured. Cell viability was measured using the NRU assay. For comparison purposes cytotoxicity curves were expressed either as a function of puff number or nicotine released in the basal media of the exposure chambers. Each laboratory performed 1 exposure of two different Transwell® at the same time (five independent replicates) for 1R6F and ENDS. The negative controls were: a “AIR” control (cells exposed to CFP filtered laboratory air at the same puffing regime during exposure run); a submerged incubator control (INC) and an “ALI” incubator control. The positive control consisted of cells exposed to 1 ml of basal and 0.5 ml of apical sodium dodecyl sulphate at 350 μM.

**Fig. 2.**
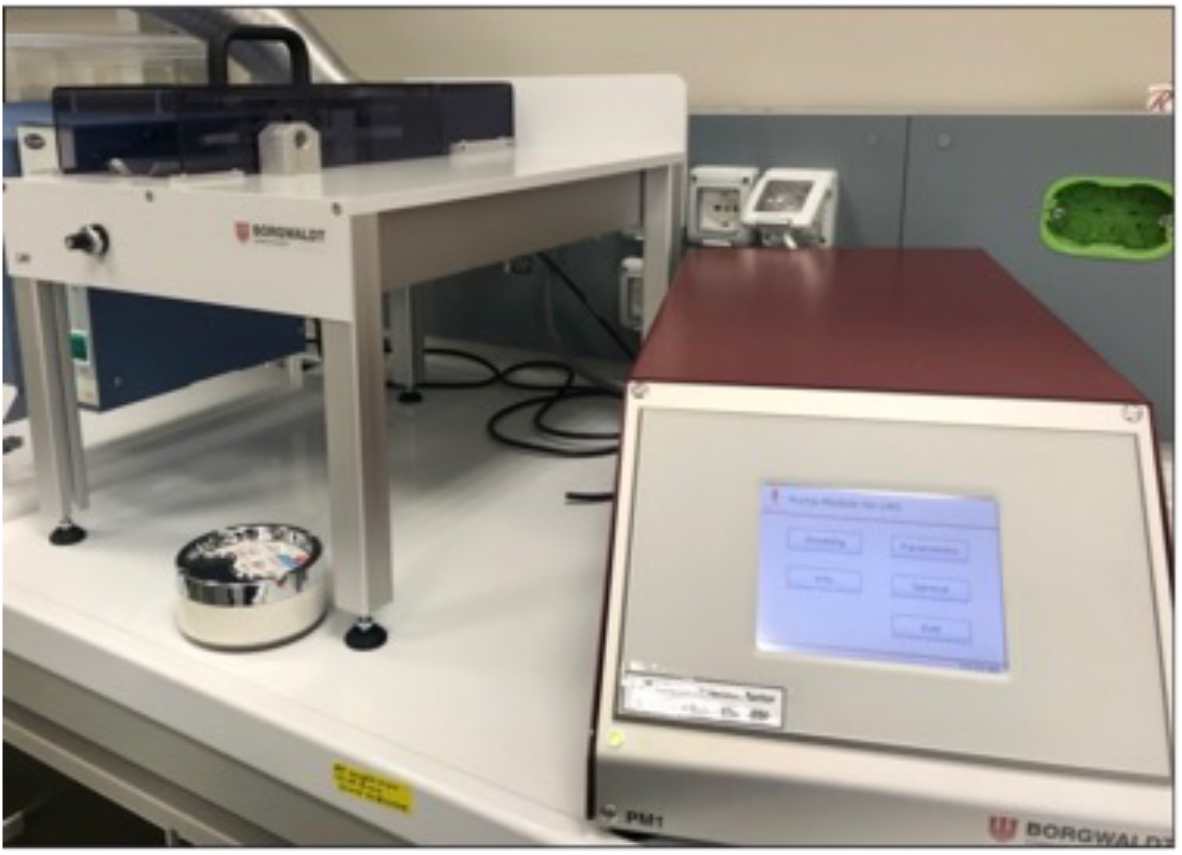
Borgwaldt LM1 smoking machine

### Cytotoxicity Test

As per replicated papers, we selected the Neutral Red Uptake (NRU) cytotoxicity assay as a benchmark assay to evaluate the cytotoxic effects of smoke (WS and VP) and aerosol on bronchial epithelial cells *in vitro*. NRU is the most recognized assay for cytotoxicity evaluation in the context of tobacco products testing (Belushkin et al., 2014), and it is also included in the first non-genotoxicity *in vitro* assay accepted for the regulatory evaluation of chemical compounds (European Commission, 2000; OECD/OCDE, 2004; Repetto et al., 2008). The NRU cytotoxicity test is a cell viability chemosensitivity test based on the ability of viable cells to incorporate and bind Neutral Red (NR), a supra-vital cationic dye that spontaneously penetrates cell membranes by non-ionic diffusion and accumulates in lysosomes. Alterations of biological membranes leading to lysosomal fragility can result in a decrease in the absorption and binding of NR. This makes it possible to distinguish between viable, damaged or dead cells (Putnam et al., 2002). After 24 h recovery period, UltraCULTURE™ was removed and stored at −80 °C for the cytokine evaluations. The exposed NCI-H292 cells were washed twice with phosphate buffered saline (PBS). Then, cells were incubated with NR dye (0.05 g/L in UltraCULTURE™) for 3 h at 37 °C, 5% CO_2_ in a humidified atmosphere. Next, NCI-H292 cells were washed with PBS to remove unincorporated dye. The incorporated NR was eluted from NCI-H292 cells by adding 500 μl of destain solution (50% ethanol, 49% distilled water, 1% glacial acetic acid; V:V:V) to each insert, and incubated for 10 min at 300 rpm on a plate shaker. NR extracts were transferred to a 96-well plate in duplicate (100 μl per well). The optical density of NR extracts was read with a microplate spectrophotometer at 540 nm using a reference filter of 630 nm. Blank insert (without cells) was used to assess how much NR solution stains the Transwells® membranes. Background measurement from Blank was subtracted from each measurement.

### Inflammatory and tissue remodeling mediator secretion

Commercially available ELISA kits were used to measure Interleukin-6 (IL-6 human Instant Elisa™ kit; Catalog Number BMS213INST), Interleukin-8 (IL-8 human Instant Elisa™ kit; Catalog Number BMS204-3INST) and MMP-1 (MMP1 Human ELISA Kit; Catalog Number EHMMP1) concentration. The assays were performed according to the manufacturer’s protocol. Absorbance was measured at a wavelength of 450 nm, and biomarker concentration was calculated from a standard curve generated with purified proteins. The detection limits as specified by the manufacturer were 0.92 pg/mL (IL-6), 1.3 pg/mL (IL-8), 8.0 pg/mL (MMP-1). Each measurement was performed in duplicate. These three cytokines released from cells 24 hours after the exposure to cigarette smoke were evaluated only by comparing between the effects of WS and VP stimulation under the two regimes (ISO and HCI).

### Statistical analysis

All raw data produced by each centre were tabulated and processed using Excel software (Microsoft, Redmond, WA, USA). Interlaboratory Studies (ILS) R package was used to assess consistency of results. Particularly, the repeatability deviation (Sr), the deviation between the means of laboratories (SB), and the reproducibility deviation (SR) are estimated. NRU cell viability data of 1R6F dose response curves were also assessed for reproducibility between labs by linear regression analysis of NRU cell viability (expressed as percentage of AIR control) obtained from each center. Moreover, “Bland and Altman” plots were created to describe the level of agreement between the different centers. GraphPad Prism 8 software was used to determine the IC50 values for each exposure (ISO WS, ISO VP, HCI WS, and HCI VP) by fitting a sigmoidal dose-response curve with a variable slope and determine the best fit values for the log IC50 of 8 parameter nonlinear regression model, and comparison between slopes. Moreover, linear regression analyses were performed to evaluate the best-fit slope between puff numbers and nicotine concentration or TPM followed by comparison between ISO and HCI slopes. Shapiro-Wilk test was performed to assess the data distribution. Inflammatory mediator data were checked by excluding values out of each ELISA standard curve. Moreover, outlier values were detected by GRUBBS’ test. Comparison of inflammatory mediator data was carried out by ANOVA with post-hoc Tukey adjustment. Comparison of NRU cell viability after exposure to 1R6F whole smoke, IQOS, GLO™, ePen and eStick aerosol was performed by using Kruskal–Wallis test followed by Wilcoxon multiple-comparison analysis with Holm’s correction. All analyses were considered significant with a p value < 0.05. R version 3.4.3 (2017-11-30) was used for data analysis and generation of graphs, otherwise stated.

## RESULTS

### Laboratory performances for 1R6F dose-response curves

We evaluated laboratory performance of NRU cell viability results for each 1R6F exposure conditions (ISO WS, ISO VP, HCI WS, and HCI VP). Significant variability was observed for D and E laboratories in performing ISO WS exposure compared to other laboratory results (**Fig. 3A**). Great interlaboratory (SR and SB) variability was observed for all the ISO WS exposure conditions (**Fig. 3B)**. Linear regression analyses of ISO WS dose response curve results showed good reproducibility between LAB-A and LAB-B (r= 0.871; p= 0.002), LAB-A and LAB-C (r= 0.774; p= 0.009), LAB-C and LAB-B (r= 0.67; p= 0.013). LAB-D and LAB-E results did not show significant correlation with the other laboratory results. Laboratory D showed more variability in performing ISO VP exposure compared to other laboratory results (**Fig. 3C**). Great interlaboratory variability (SR and SB) was observed for 2, 5, 10, 12, and 25 puff ISO VP exposure conditions. Moreover, intralaboratory variability (Sr) was observed for 2 and 5 puff ISO VP exposure conditions (**Fig. 3D)**. Linear regression analyses of ISO VP dose response curve results showed good reproducibility between LAB-A and LAB-B (R= 0.917; p= < 0.001), LAB-A and LAB-C (R= 0.946; p= < 0.0001), LAB-A and LAB-E (R= 0.715; p= 0.008), LAB-C and LAB-B (R= 0.889; p= < 0.001), LAB-C and LAB-E (R= 0.69; p= 0.011), LAB-B and LAB-E (R= 0.587; p= 0.027). LAB-D results did not show significant correlation with the other laboratory results. Variability was observed only for 2 and 4 puff HCI WS exposure of LAB-D laboratory (**Fig. 3E)**. Great interlaboratory (SR and SB) variability was observed for 2, 4, 5, 6, and 8 HCI WS puff exposures (**Fig. 3F)**. Linear regression analyses of HCI WS dose response curve results showed good reproducibility between LAB-A and LAB-B (R= 0.77; p= 0.004), LAB-A and LAB-C (R= 0.586; p= 0.027), LAB-A and LAB-D (R= 0.729; p= 0.007) LAB-A and LAB-E (R= 0.763; p= 0.005), LAB-C and LAB-E (R= 0.624; p= 0.02), LAB-B and LAB-E (R= 0.677; p= 0.012), LAB-B and LAB-D (R= 0.745; p= 0.006), LAB-D and LAB-E (R= 0.636; p= 0.018). More variability was observed for LAB-C results in performing HCI VP exposure compared to other laboratory results (**Fig. 3G)**. Greater interlaboratory variability (SR and SB) was observed for 8, 10, and 15 puff HCI VP exposure conditions (**Fig. 3H)**. Linear regression analyses of HCI VP dose response curve results showed good reproducibility between LAB-A and LAB-B (R= 0.861; p= 0.003), LAB-A and LAB-D (R= 0.8557; p= 0.003) LAB-A and LAB-E (R= 0.671; p= 0.024), LAB-B and LAB-E (R= 0.663; p= 0.014), LAB-B and LAB-D (R= 0.94; p< 0.0001), LAB-B and LAB-E (R= 0.714; p= 0.008), LAB-D and LAB-E (R= 0.702; p= 0.009).

**Figure 3.**
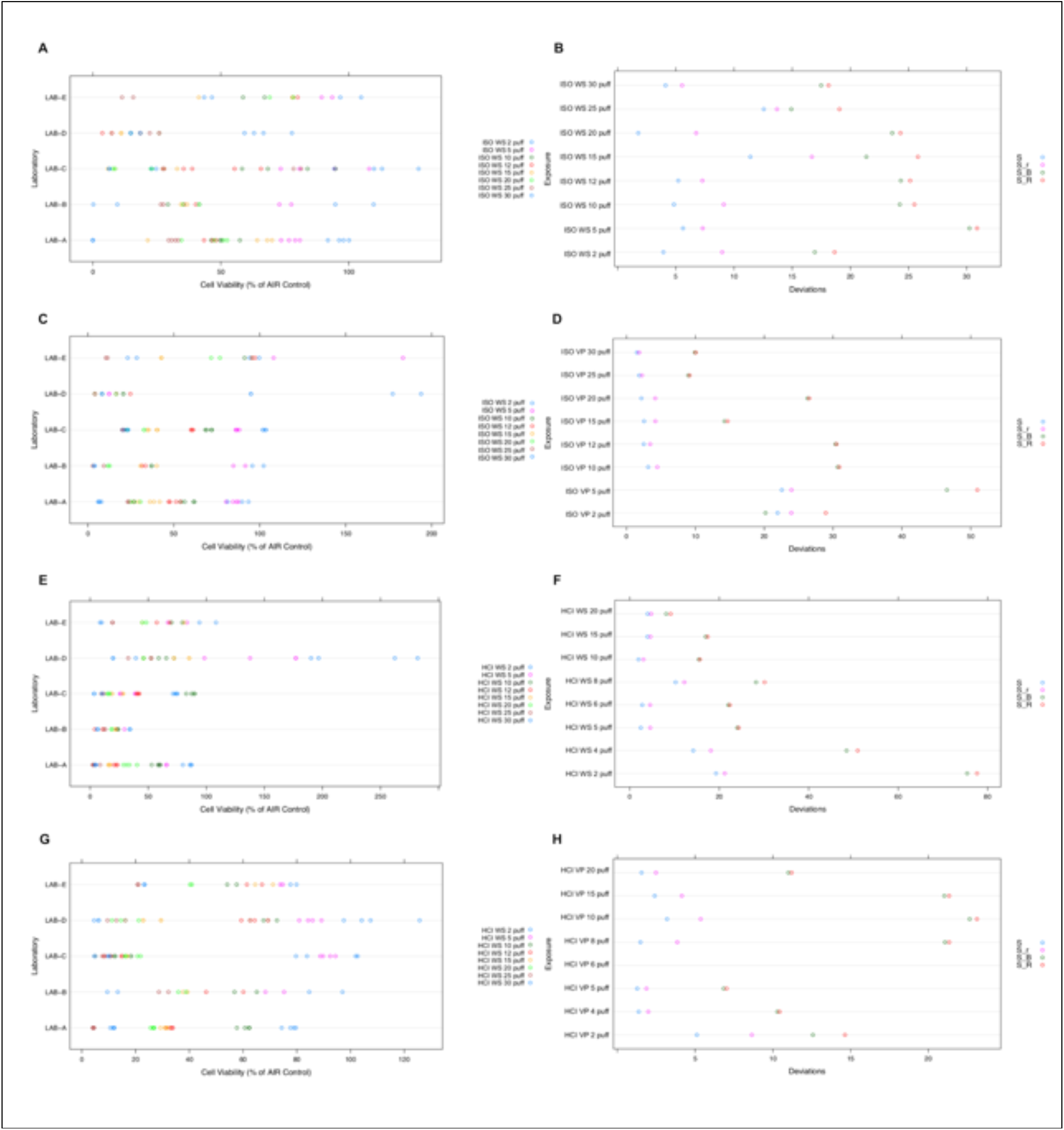
Laboratory performances for 1R6F dose-response curves following ISO and HCI regimes. ISO WS measurements of NRU cell viability (**A**) and measures of variability for each puff number (**B**); ISO VP measurements of NRU cell viability (**C**) and measures of variability for each puff number (**D**); HCI WS measurements of NRU cell viability (**E**) and measures of variability for each puff number (**F**); HCI VP measurements of NRU cell viability (**G**) and measures of variability for each puff number (**H**). S: global deviation of all laboratories, Sr: repeatability’s deviation (intra-laboratory), SB: deviation between the means of the laboratories, SR: reproducibility’s deviation (interlaboratory). WS: Whole smoke, VP: Vapour phase.

### Nicotine dosimetry

Nicotine concentration was quantified in the cell culture media after each exposure only for the experiments performed by LAB-A. Instead, the Total Particulate Matter (TPM) was quantified by LAB-A, LAB-C, and LAB-D. Dosages of nicotine by VP exposure were all under the limit of quantification (LOQ) because CFPs retain almost all nicotine released by cigarette smoke. Only WS exposure released a quantity of nicotine detectable in the exposed culture media. We observed an increase of nicotine with the increase of puff number (**Fig.4A**), and this escalation was more accentuated for the HCI regime compared to ISO regime (p= 0.0005). A similar trend was observed when the TPM was presented against puff number exposure. The TPM release under HCI regime was significantly increased compared to ISO regime with a p value < 0.0001 (**Fig. 4B**).

**Figure 4.**
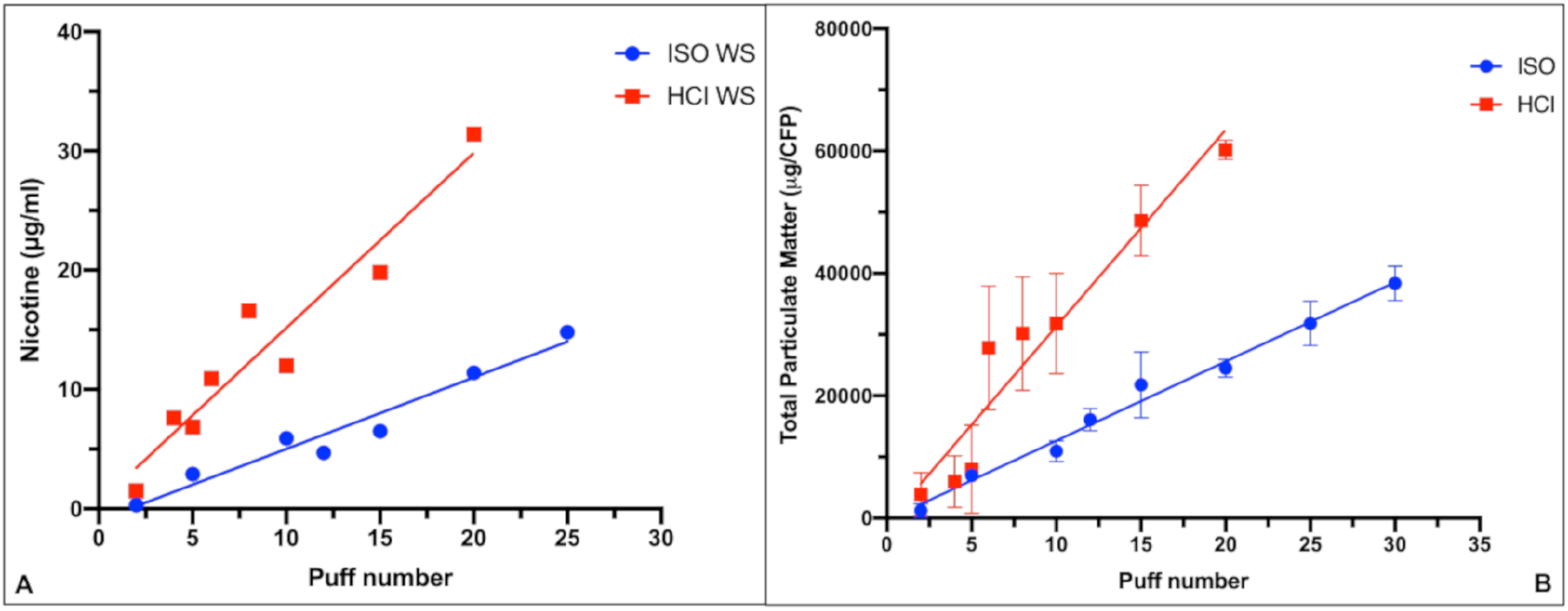
(**A**) Nicotine (μg/ml) released by each puff number exposure for ISO and HCI generated 1R6F WS. There was a statistical difference in nicotine release between ISO and HCI exposure (p= 0.0005). (**B**) Total Particulate Matter (μg/CFP) per each puff number exposure for ISO and HCI generated by 1R6F WS. There was a statistical difference in nicotine release between ISO and HCI exposure (p< 0.0001). WS: Whole smoke, VP: Vapor phase.

Using nicotine dosimetry data of WS exposure, cell viability of the biological responses was presented against exposed nicotine in the cell media for each puff number. ISO WS exposure showed an IC50 of 4.18 μg/ml of nicotine that was about the nicotine released by 10 puffs. HCI WS exposure showed an IC50 of 9.7 μg/ml of nicotine that was about the nicotine released by 5 puffs.

Based on nicotine released at IC50 dose by WS under the HCI regime, we chose to conduct the second and third phase of this study using the number of puffs able to reach the same amount of nicotine across all products (1R6F, IQOS, GLO™, ePen, and eStick). We observed that the means ± SD of released nicotine were 8.55 ± 0.78 μg/ml for 5 puffs of 1R6F HCI WS, 9.03 ± 1.31 μg/ml for 7 puffs of IQOS, 8.7 ± 1.3 μg/ml for 8 puffs of GLO™, 8.47 ± 1.54 μg/ml for 10 puffs of ePen, and 8.8 ± 2.4 μg/ml for 25 puffs of eStick.

### Effect of WS and VP ISO regime on H292 cell viability and inflammatory mediators

Based on laboratory performance results, LAB-D and LAB-E NRU data of ISO WS exposure were excluded from ISO WS IC50 determination and consequently from inflammatory mediator analyses. Also, LAB-D data of ISO VP exposure were excluded from ISO VP IC50 determination and consequently from inflammatory mediator analyses. LAB-E provided inconsistent inflammatory mediator results, which were excluded from analyses. Following the ISO regime, 1R6F WS decreased cell viability from 2 puffs to 30 puffs with an IC50 value of 10.47 puffs. Also, VP exposure decreased cell viability from 2 puffs to 30 puffs, but with an IC50 value of 11.76 puffs. The IC50 for WS exposure was reduced by about 12 % than that required following exposure to VP (**Fig. 6**), but no significant difference was observed between the two IC50 values (p= 0.098).

**Figure 5.**
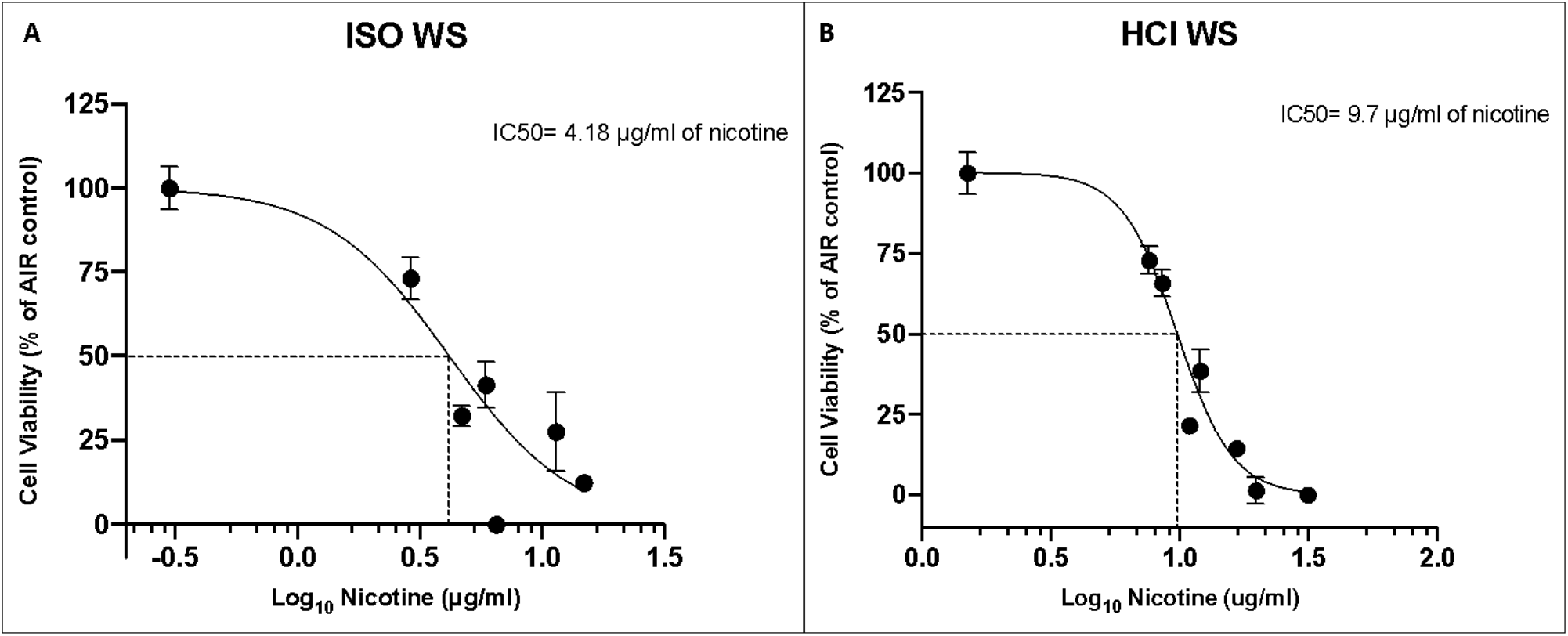
Neutral Red Uptake (NRU) cell viability of NCI-H292 bronchial epithelial cells exposed to 1R6F ISO WS **(A**) and HCI WS (**B**). Data points are mean ± SD from 4 replicates (LAB-A). Biological response data are presented as a function of nicotine concentration in the exposed media. The IC50 of ISO WS was 4.18 μg/ml of nicotine (Log_10_= 0.62); the IC50 for HCI WS was 9.7 μg/ml of nicotine (Log_10_= 0.99). WS: Whole smoke, VP: Vapor phase.

**Figure 6.**
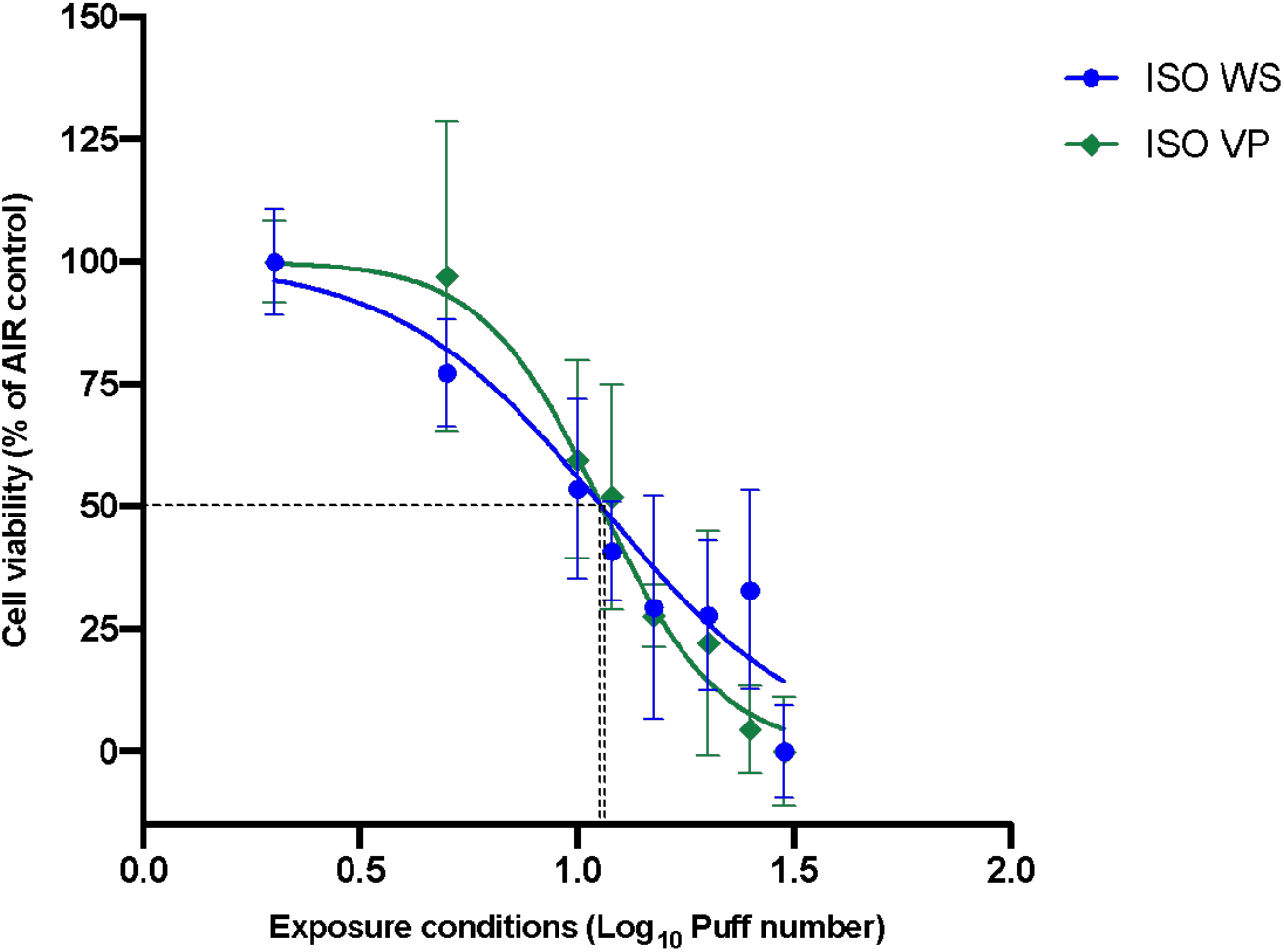
Cytotoxicity of NCI-H292 cells exposed to 1R6F ISO WS 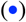 and ISO VP 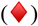. Data points are mean ± SD from ten replicates. There was no significant (p= 0.098) increase in IC50 for VP (11.76 puffs; Log_10_=1.07) exposure compared to IC50 for WS exposure (10.47 puffs; Log_10_=1.02).

Following exposure to a range of both ISO WS and VP puff numbers, all the inflammatory mediators showed highest values for the lower puff numbers with a decrease as the puff number increased (**Fig. 7A–9A**). Also, no significant differences (p values > 0.05) were observed between WS and VP exposures for all the inflammatory mediators. IL-6 concentrations at the highest puff numbers of VP and WS were significantly (p < 0.05) lower than those seen in the AIR control (**Fig. 7A**). Only IL8 inflammatory mediator released at 2 and 5 puffs of WS exposure were significantly increased (p< 0.001) when compared to the AIR control (**Fig. 8A**). When IL6 and IL8 concentrations were normalized to NRU cell viability, we observed an increase of these concentrations at higher puff numbers (**Fig. 7B and 8B**). Moreover, IL6 and IL8 concentrations following ISO WS exposure seems to be higher than VP exposure, but no significant differences were observed. No significant variations were observed when NRU cell viability normalization was applied to MMP1 results (**Fig. 9B**).

**Figure 7.**
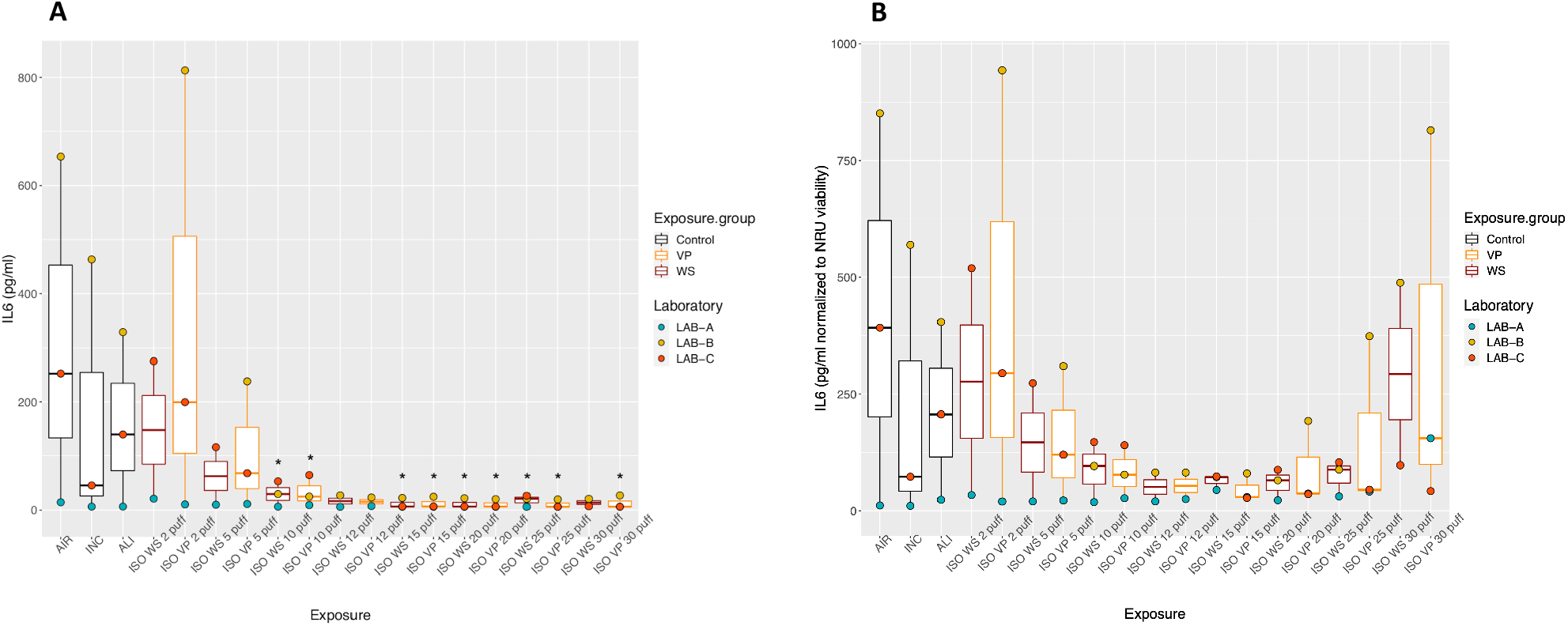
IL6 inflammatory mediator secretion following ISO WS and VP exposures. Boxplots represent “minimum”, first quartile (Q1), median, third quartile (Q3), and “maximum” of IL6 concentrations (**A**) and IL6 concentration-normalized to NRU viability (**B**) for each exposure condition. The laboratory color coded points represent the mean of each laboratory results. AIR: Air control, INC: Incubator control, ALI: Air–liquid interface control, WS: Whole smoke, VP: Vapour phase. (*) p < 0.05 compared to AIR control.

**Figure 8.**
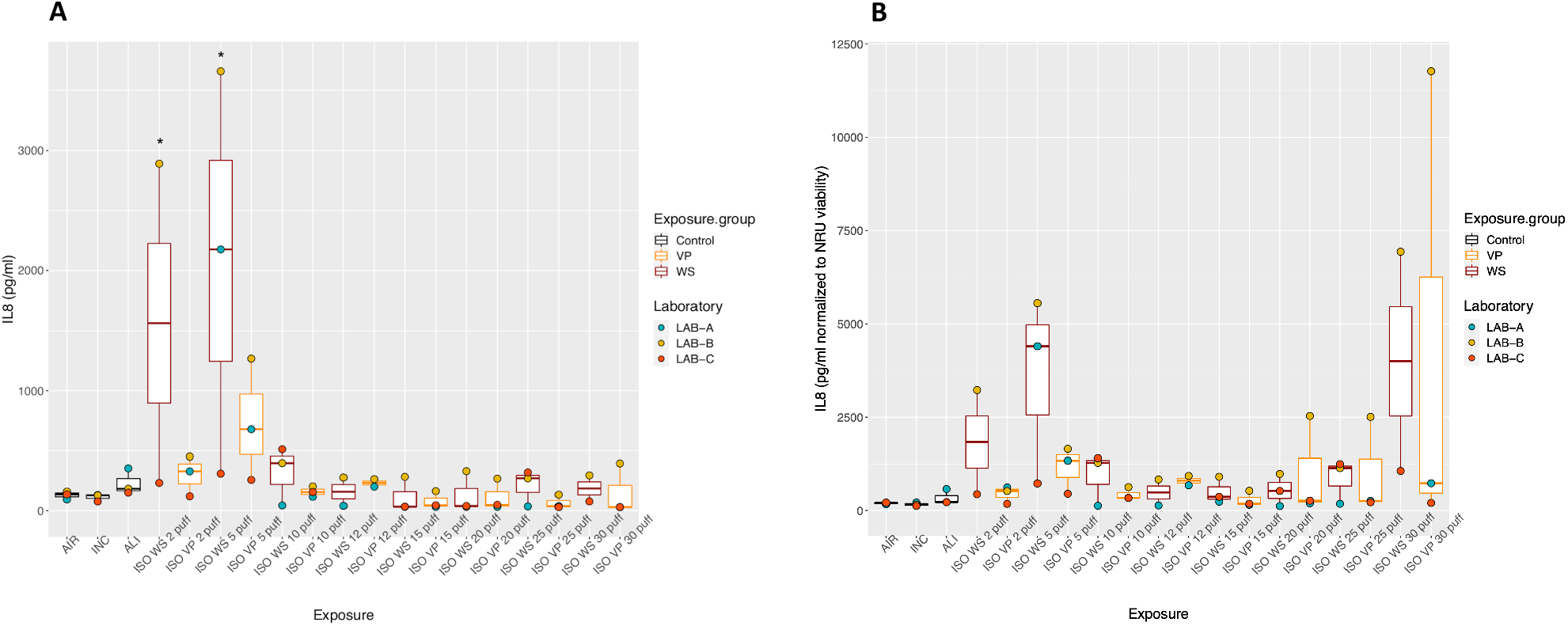
IL8 inflammatory mediator secretion following ISO WS and VP exposures. Boxplots represent “minimum”, first quartile (Q1), median, third quartile (Q3), and “maximum” of IL8 concentration (**A**) and IL8 concentration-normalized to NRU viability (**B**) for each exposure condition. The laboratory color coded points represent the mean of each laboratory results. AIR: Air control, INC: Incubator control, ALI: Air–liquid interface control, WS: Whole smoke, VP: Vapour phase. (*) p < 0.05 compared to AIR control.

**Figure 9.**
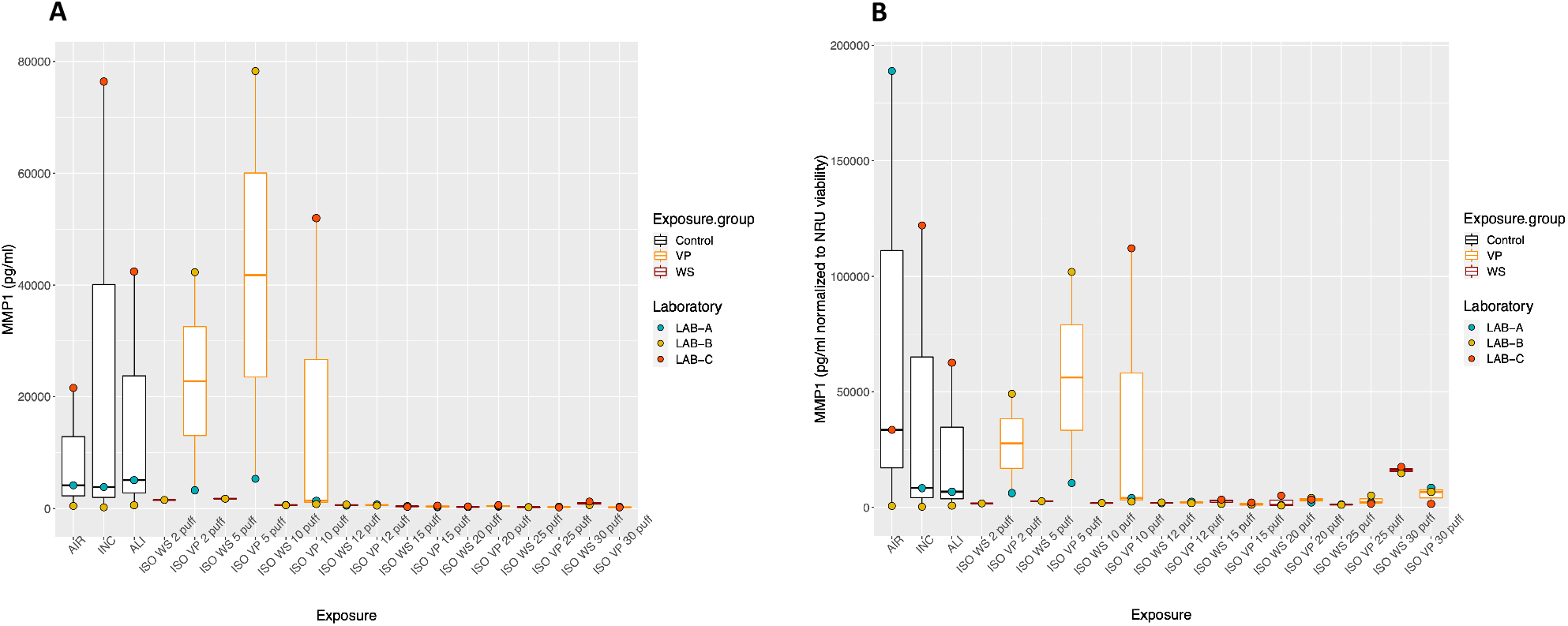
MMP1 tissue remodeling mediator secretion following ISO WS and VP exposures. Boxplots represent “minimum”, first quartile (Q1), median, third quartile (Q3), and “maximum” of MMP1 concentration (**A**) and MMP1 concentration-normalized to NRU viability (**B**) for each exposure condition. The laboratory color points represent the mean of each laboratory result. AIR: Air control, INC: Incubator control, ALI: Air–liquid interface control, WS: Whole smoke, VP: Vapor phase.

### Effect of WS and VP HCI regime on H292 cell viability and inflammatory mediators

Based on laboratory performance results, LAB-C NRU data of HCI VP exposure were excluded from HCI VP IC50 determination and consequently from inflammatory mediator analyses. LAB-E provided inconsistent inflammatory mediator results, which were excluded from analyses. Following the HCI regime 1R6F whole smoke decreased cell viability from 2 puffs to 20 puffs with an IC50 value of 5.14 puffs. Also, VP exposure decreased cell viability from 2 puffs to 20 puffs but with an increased IC50 value of 6.22 puffs. The IC50 for WS exposure was significantly reduced (p= 0.046) of about 21 % than that required following exposure to VP (**Fig. 10**).

**Figure 10.**
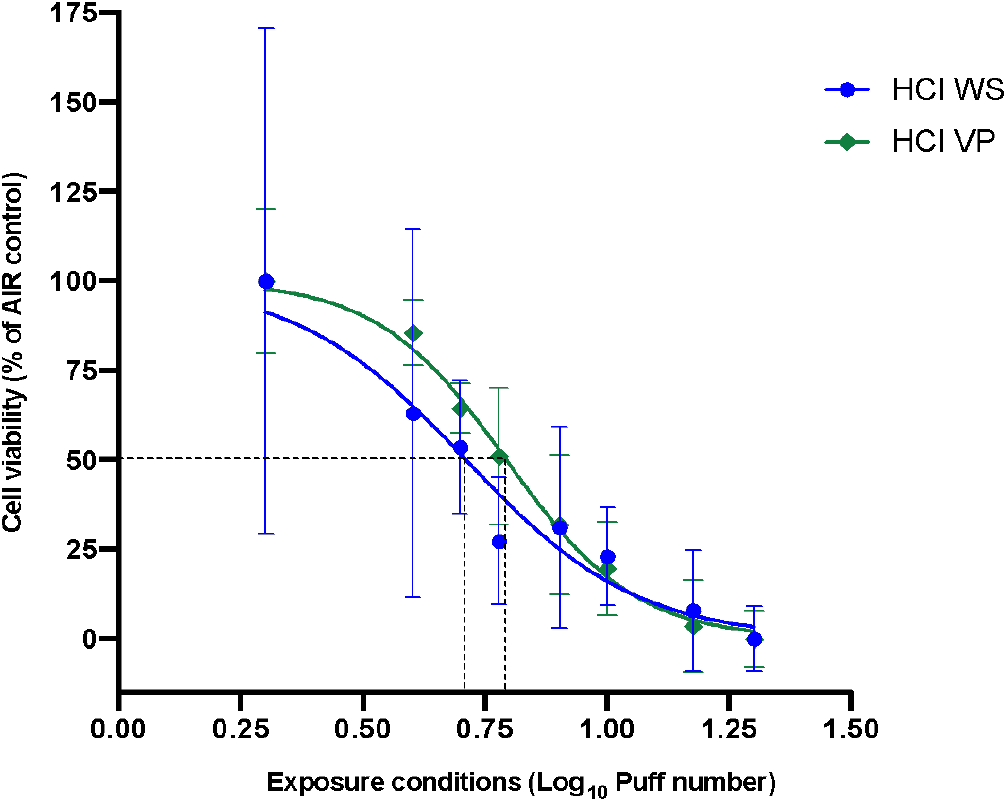
Cytotoxicity of NCI-H292 cells exposed to 1R6F HCI WS 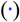 and HCI VP 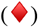. Data points are mean ± SD from 16 replicates for WS and 10 replicates for VP. Significant (p= 0.046) increase in IC50 for VP (6.22 puffs; Log_10_= 0.79) exposure compared to IC50 for WS (5.14 puffs; Log_10_= 0.71) exposure.

Following exposure to a range of both HCI WS and VP puff numbers, IL6, IL8, and MMP1 mediators showed highest values for the lower puff numbers with a decrease as the puff number increases (**Fig. 11A–13A**). Also, VP exposures seem to increase the release of IL6, IL8 and MMP1 mediators, but no significant differences (p values > 0.05) were observed between WS and VP exposures for all the inflammatory mediators. IL-6 concentrations at the highest puff numbers of VP and WS were significantly (p < 0.05) lower than those seen in the AIR control (**Fig. 11A**). IL8 inflammatory mediator released at 2 puffs of VP exposure was significantly increased (p< 0.001) when compared to the AIR control (**Fig. 12A**). When IL6 and IL8 concentrations were normalized to NRU cell viability, we observed an increase of these concentrations at higher puff numbers (**Fig. 11B and 12B**). Slight variations were observed when NRU cell viability normalization was applied to MMP1 results (**Fig. 13B**).

**Figure 11.**
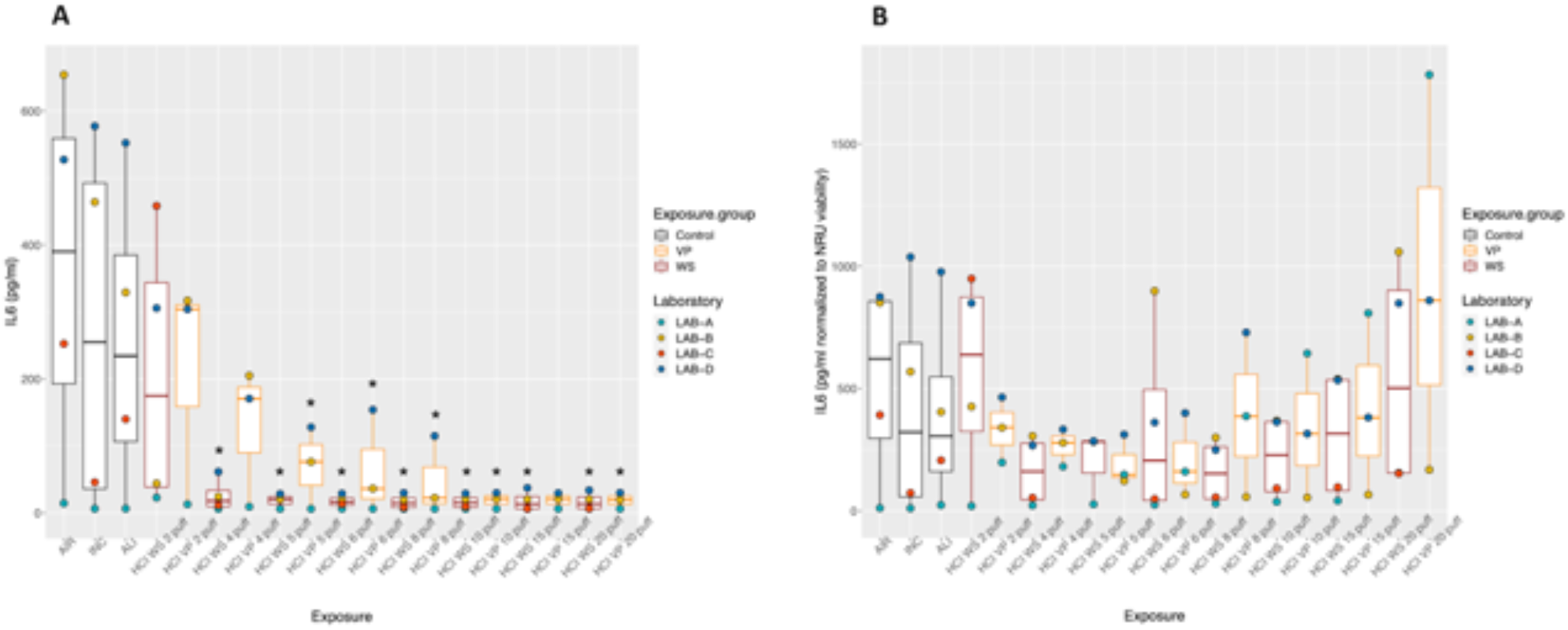
IL6 inflammatory mediator secretion following HCI WS and VP exposures. Boxplots represent “minimum”, first quartile (Q1), median, third quartile (Q3), and “maximum” of IL6 concentrations (**A**) and IL6 concentration-normalized to NRU viability (**B**) for each exposure condition. The laboratory color coded points represent the mean of each laboratory result. AIR: Air control, INC: Incubator control, ALI: Air–liquid interface control, WS: Whole smoke, VP: Vapour phase. (*) p < 0.05 compared to AIR control.

**Figure 12.**
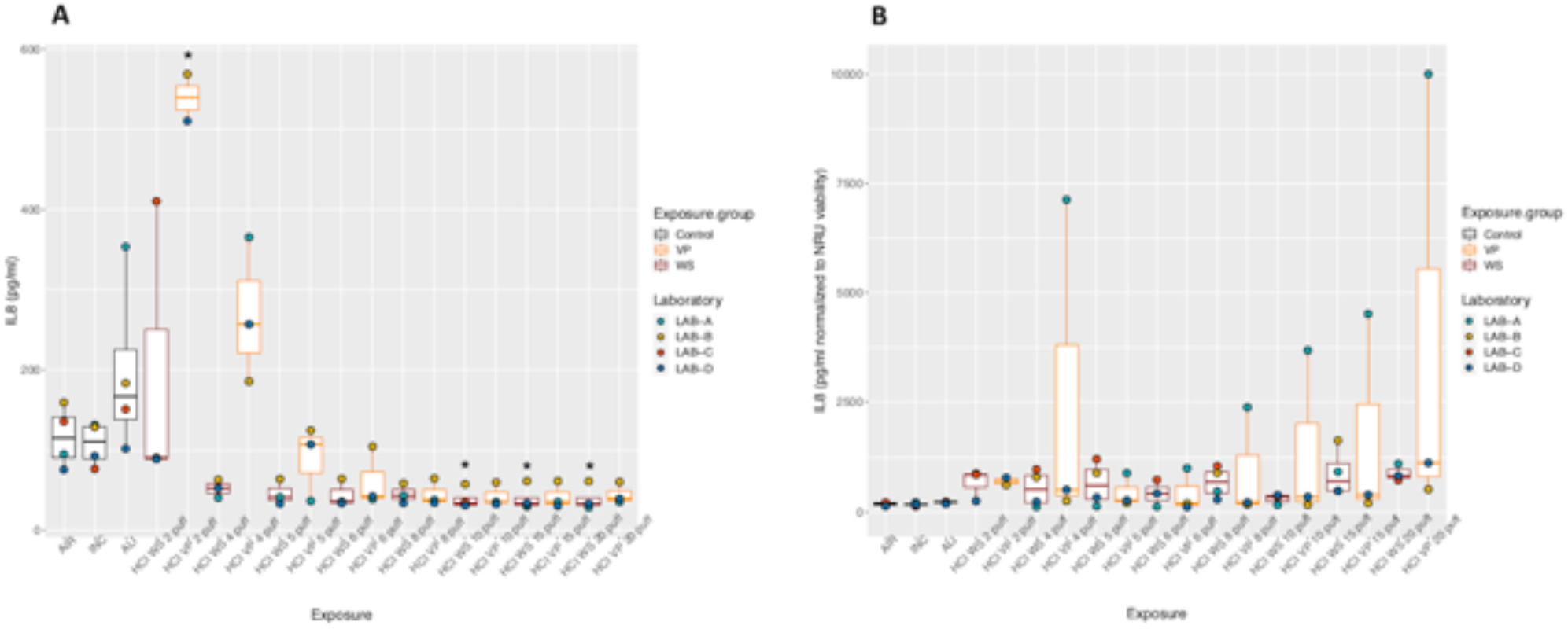
IL8 inflammatory mediator secretion following HCI WS and VP exposures. Boxplots represent “minimum”, first quartile (Q1), median, third quartile (Q3), and “maximum” of IL8 concentration (**A**) and IL8 concentration-normalized to NRU viability (**B**) for each exposure condition. The laboratory color coded points represent the mean of each laboratory result. AIR: Air control, INC: Incubator control, ALI: Air–liquid interface control, WS: Whole smoke, VP: Vapour phase. (*) p < 0.05 compared to AIR control.

**Figure 13.**
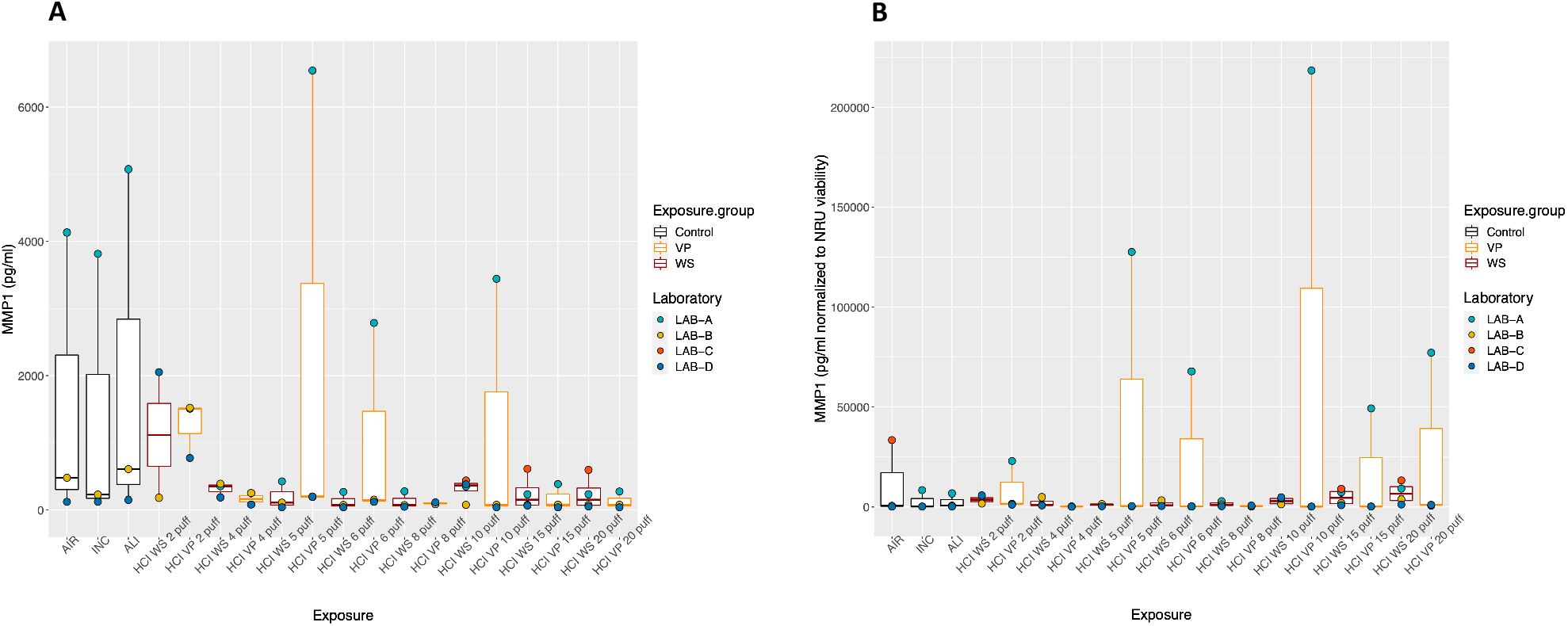
MMP1 tissue remodeling mediator secretion following HCI WS and VP exposures. Boxplots represent “minimum”, first quartile (Q1), median, third quartile (Q3), and “maximum” of MMP1 concentration (**A**) and MMP1 concentration-normalized to NRU viability (**B**) for each exposure condition. The laboratory color coded points represent the mean of each laboratory result. AIR: Air control, INC: Incubator control, ALI: Air–liquid interface control, WS: Whole smoke, VP: Vapour phase.

### Laboratory performances for exposure to 1R6F, THPs, and e-Cigs comparison

We evaluated laboratory performance for each exposure condition (1R6F 5 puffs, IQOS). More variability was observed for 1R6F 5 puffs exposure among all laboratories (**Fig. 14A**). Indeed, higher interlaboratory (SR and SB values) variability was observed for 1R6F exposure condition compared to other products (**Fig. 14B)**.

**Figure 14.**
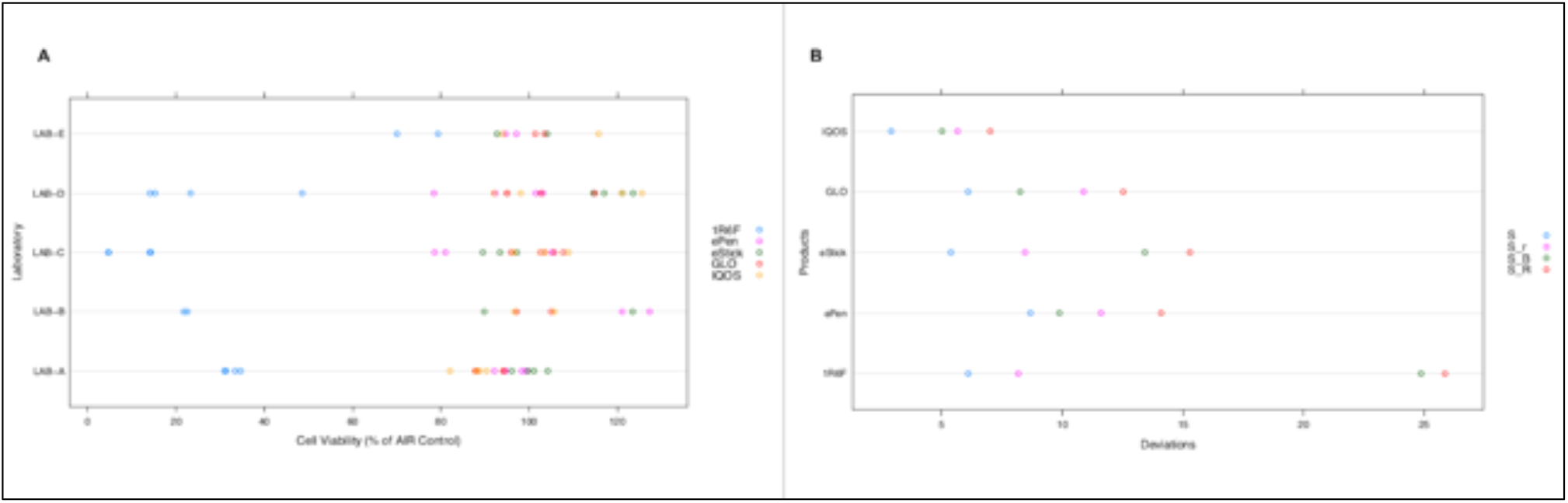
Laboratory performances for exposure to 1R6F, THPs, and e-Cigs comparison. Measurements of NRU cell viability (**A**) and measures of variability for each test product (**B**); S: global deviation of all laboratories, Sr: repeatability deviation, SB: deviation between the means of the laboratories, SR: reproducibility deviation.

### Effect of THPs and e-Cig exposure on H292 cell viability compared to 1R6F exposure

Comparison of NRU cell viability among all product exposures showed a significant difference with an overall p value< 0.0001 (**Fig. 15**). Particularly, we observed a significant reduction of cell viability after exposure to 1R6F smoke, 26.45 % (14.5-33.1), compared to AIR control (p= 0.009). No reduction in cell viability was also observed in H292 cells exposed to IQOS, 93.34 % (88.2-103.1), GLO™, 95.04 % (89.6-103.3), ePen, 97.57 % (92.2-102.3), and eStick, 101.09 % (96.1-114.5), aerosol compared to AIR controls. Cross comparisons among all the tested products showed that only cell viability reduction after 1R6F exposure is significantly different compared to all the other products (p values < 0.0001).

**Figure 15.**
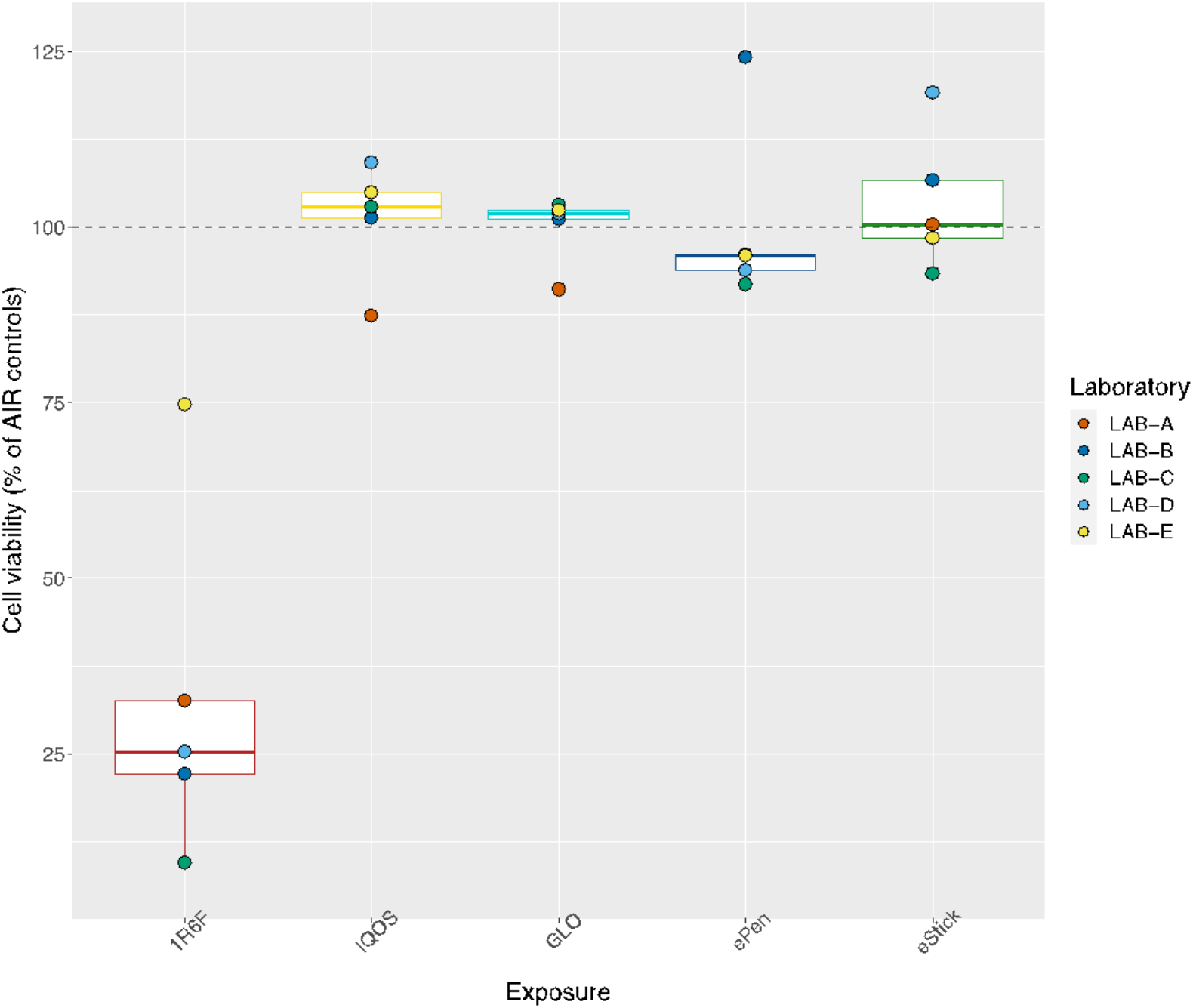
Comparison of NRU cell viability after exposure to 1R6F whole smoke, IQOS, GLO™, ePen and eStick vapours. Boxplots represent “minimum”, first quartile (Q1), median, third quartile (Q3), and “maximum” of MMP1 concentration (**A**) and MMP1 concentration-normalized to NRU viability (**B**) for each exposure condition. The laboratory color coded points represent the mean of each laboratory results. The medians (IQR range) were respectively 26.45 % (14.5-33.1) for 1R6F, 93.34 % (88.2-103.1) for IQOS, 95.04 % (89.6-103.3) for GLO™, 97.57 % (92.2-102.3) for ePen, and 101.09 % (96.1-114.5) for eStick.

## Discussion

In 2019 the Replica project was launched with the specific purpose of verifying the results obtained from the most important studies on cytotoxicity of electronic nicotine delivery systems *in vitro* from tobacco companies. Our aim was to verify the robustness and reliability of the conclusions reached by these studies, replicating their experiments as best as possible, considering the rapid evolution of this sector and of the products available on the market. The first studies were selected on the basis of specific criteria. The forward thrust of this replication study was the execution of the experiments in rounds in 5 different international laboratories.

The initial studies we have chosen for the first year were conducted on cytotoxicity and inflammation on a model easy to replicate, in order to achieve the main goal, which was to fine-tune the collaboration between the international partners of the various laboratories participating in the study. A secondary, but relevant, outcome was to harmonize protocols of exposure and cell viability studies between the partners of the study in order to produce accurate, reliable and unbiased results as much as possible. This study contributes to establish standardized procedures to test the effects *in vitro* of ENDS’ aerosol on human bronchial epithelial cells, applicable to different products available in the market in order to establish the effects of aerosol on human cells, contributing to the regulatory science of ENDS. Correlating the results obtained in this multi-centric study with those obtained by Azzopardi et al. (Azzopardi et al., 2015) we observed a difference between the cytotoxicity induced by cigarette WS and VP comparable to that highlighted in the original study, confirming that a substantial part of the acute cytotoxicity induced by cigarette smoke on the cells of the lung epithelium is mediated by the volatile component of the smoke and not by TPM or nicotine, in a large part effectively trapped by CFPs. A similar result was obtained in the comparison of the cytotoxicity induced by conventional cigarette smoke and the aerosol of electronic cigarettes, which for a comparable amount of nicotine released in the culture medium of the exposure chamber appears to be non-toxic to lung cells, as corroborated by the assessment in the second paper replicated (Azzopardi et al., 2016). Again, a comparable result is found between the cytotoxicity induced by cigarette smoke and the aerosol of THPs, since both reduced risk products (GLO™ and IQOS) seem to be extremely less toxic compared to cigarette smoke, producing a cytotoxic effect of about 3.5% (mean) compared to 73.55 % induced by the former.

In this study, compared to the replicated studies, we added a direct comparison between these different ENDS, comparing e-cigs to THP which, with the settings used and at these quantities of nicotine released, seem equally unable to induce cytotoxicity on bronchial epithelial cells.

It is well-known that tobacco smoke is able to induce an imbalance between oxidants and antioxidants in the airways leading to oxidative stress (Emma et al., 2018), increased mucosal inflammation, and increased expression of inflammatory cytokines (such as IL-8, IL-6 and TNF-alfa) (Strzelak et al., 2018). Azzopardi and colleagues (Azzopardi et al., 2015) found that AIR exposure increased 3-4 times IL-6 and IL-8 production compared to ALI and INC conditions, whereas we found that AIR exposure increased only 2 times the production of the inflammatory cytokines. Moreover, in the original paper by Azzopardi and colleagues, exposure of NCI-H292 cells to smoke increased the quantity of IL-6 and IL-8 released from cells, in particular exposure to WS rather than VP. The quantity of IL-6 released from bronchial cells exposed to WS was increased compared to AIR exposure, but decreased in cells exposed to VP, leading to the conclusion that the proinflammatory stimulus was mainly due to the particulate components of the smoke (Azzopardi et al., 2015). Regarding the inflammatory (IL-6 and IL-8) mediators the results obtained in our study regarding the effect of cigarette WS and VP are quite different because we observed a significant dose-dependent decrease of IL-6 release from bronchial epithelial cells exposed to WS compared to AIR exposure, both with ISO and HCI regime. Notably, and contrary to the results of Azzopardi and colleagues, we observed a greater production of IL-6 from the cells exposed to VP compared to WS with the two regimes (ISO and HCI), however always lower than the exposure to AIR. Normalizing the quantity of IL-6 released with the value of viable cells by NRU assay, we observed an increase in IL-6 production which overtake the AIR exposure only by HCI regime for the highest number of puffs (20 puffs) and only by VP exposure. Unexpectedly, our results on IL-6 seem to indicate an anti-inflammatory effect of smoke, but normalizing data to viable cells, it seems to be clear that these cells need a more consistent or prolonged stimulus to activate the release of this cytokine (**Fig. 11**). Probably, direct exposure of cells to undiluted smoke has an exacerbating effect on bronchial cells, generating strong cytotoxicity and initially deactivating the cellular machinery that is unable to respond promptly to the insult represented by smoking. In more prolonged and extreme conditions of exposure (HCI, 20 puffs), the proinflammatory response also seems to appear, but it seems to be greatly dependent from volatile components.

For IL-8 we observed no significant difference between the three controls (ALI, INC and AIR), but we observed a significant increase in cytokine production by NCI-H292 cells exposed to 2 and 5 puffs of WS under the ISO regime and to 2 puffs of VP by HCI regime. Increasing the number of puffs for both the regimes we observed a substantial decrease in IL-8 release. Exposure to WS was more effective than VP in inducing IL-8 release under the ISO regime, but there was a reverted effect under the HCI regime, since the exposure to VP was more effective than to WS. Also, in this case the data were somewhat different when normalizing for cell viability, in particular for the higher number of puffs both under ISO (30 puffs) and HCI regimen (20 puffs). Finally, Azzopardi et al. (2015) showed no significant differences between INC, ALI and AIR control in MMP-1 release from bronchial cells and an increased MMP-1 release from cells exposed to WS and VP particularly at high smoke dilution with air and mainly induced by WS. In our study we observed no differences between ALI, INC and AIR controls, as previously reported (Azzopardi et al., 2015). Moreover, we observed a slight, but no significant, increase in MMP-1 release in NCI-H292 bronchial cells exposed to VP under ISO regime and only at 2 and 5 puffs, and an absolutely irrelevant release of MMP-1 from cells exposed to WS under both ISO and HCI regimes. Cigarette smoke is known to induce MMP-1 mRNA and protein expression in human airway cells (Mercer et al., 2004), consequently inducing excessive matrix remodeling in smokers leading to emphysema. Also in this case, it seems that in our model of exposure the impact of smoking is too strong and in fact as the number of puffs increases the cells seem unable to respond to the damage induced by smoking. Probably, to study active responses of cells to smoking, direct exposure to undiluted smoke could be too extreme, given that smoking is already well-known for its cytotoxic activity on human cells. Therefore, a milder model of exposure, as done by Azzopardi and colleagues, to smoke diluted with air, could be a better approach to evaluate the active responses of cells.

The reproducibility of the study by Azzopardi et al. (2015) has been assessed for cytotoxicity. They found that the EC50 for WS exposure was 1:54 (smoke:air, vol:vol) and 1:46 for VP exposure alone (p < 0.005) using the ISO regime, so that the average difference in cell viability between the two dose response curves was 11%, and thus VP constituted 89% of the total toxicity of WS by ISO regime. In our study the IC50 for WS exposure was reduced of about 12 % than that required following exposure to VP for ISO regime (**Fig. 6**), and the IC50 for WS exposure was reduced of about 21 % (p= 0.046) than that required following exposure to VP under HCI regime (**Fig. 10**). This proves that VP constitutes 88% of the total toxicity of WS under ISO regime and 79% under HCI regime. Despite the difference in cigarette used (3R4F *vs*. 1R6F) and method of exposure (air-diluted smoke *vs* undiluted smoke) our study confirms that most of the cigarette smoke-induced cytotoxic damage on bronchial epithelial cells is due to the substances contained in the VP of the smoke. Instead, the data of Azzopardi et al. on the comparison of TPM production between cigarettes smoked with the ISO and HCI regimes was not confirmed by our study, since they reported no significant effect of smoking regime, whereas we observed a significant increase in the TPM produced by the cigarette smoked with the more intense HCI regime (**Fig. 4B**), as well as for the nicotine produced (**Fig. 4A**). Moreover, our results on cytotoxicity induced by the two different smoking regimes seems to be in contrast with that reported by Azzopardi and colleagues (Azzopardi et al., 2015). They reported that the EC50 values for WS derived from the ISO regimen were significantly more toxic than that derived under the HCI regimen. Moreover, they reported that TPM produced under the two regimes was substantially the same as a function of the smoke dilution value. The same comparison on undiluted smoke from our data shows that the ISO regime produces significantly less TPM and nicotine as a function of the puff number (**Fig. 4**). Moreover, comparing the IC50 obtained by exposing the cells to the two regimes in function of puffs it seems to be clear that smoke produced under the HCI regime is two-fold more toxic compared to smoke produced under ISO regime (**Fig. 5**). Notably, the IC50 under ISO regime is reached after 10 puffs, whereas IC50 under HCI regime is reached at 5 puffs. It is implied that HCI is a regimen that produces a doubled cytotoxic effect compared to ISO, even if the nicotine concentration at IC50 by ISO is 4.18 μg/ml (in 10 puffs) and instead by HCI is 9.7 μg/ml (in 5 puffs). In this evaluation it must be taken into account that, as demonstrated by both Azzopardi et al. (2015) and by this study, 88-89% of the cytotoxicity induced by cigarette smoke is caused by VP and, therefore, not by TPM. Another conclusion that we can gather by these results is that nicotine is not directly responsible for the cytotoxic effect. Jaunky and colleagues (Jaunky et al., 2018) compared the cytotoxic effects of cigarette smoke (3R4F) to aerosol from two THPs on NCI-H292 human lung epithelial cells reporting a statistically significant reduction in biological response from THPs, with >87% viability relative to 3R4F at a common aerosol dilution (1:40, aerosol:air). Therefore, we assessed the cytotoxicity of THP aerosol relative to cigarette smoke from a 1R6F cigarette exposing cells to the undiluted smoke/aerosol reaching the same nicotine amount in the media of the exposure chamber with cells. Cell viability was determined by NRU assay exposed at the ALI, as per original paper by Jaunky et al. We observed that NCI-H292 cells exposed to 5 puffs of 1R6F undiluted smoke using HCI regime had a significant reduction of viability (26.45% viable cells) whereas IQOS aerosol (7 puffs) and GLO™ PRO aerosol (8 puffs) produced under HCI regime showed respectively 93.34% and 95.04% of viable cells compared to AIR control (**Fig. 15**). These data indicate that THPs showed >67% viability relative to 1R6F with undiluted aerosol/smoke, confirming the reduced cytotoxic potential of THPs relative to a conventional cigarette. The third paper by Azzopardi and colleagues (Azzopardi et al., 2016) compared cytotoxic effects of cigarette smoke (3R4F) to aerosol from an electronic cigarette (Vype ePen) on the same cells (NCI-H292). Moreover Azzopardi et al. used another electronic device, the Vype eStick, for dosimetric assessment of their smoking machine. In regards to cytotoxicity assessment, Azzopardi observed that on an aerosol dilution basis the Vype ePen aerosol produced under a modified HCI (mHCI) regime (puff volume, duration and frequency of 55 ml, 2 s, 30 s, (55/2/30) and with a rectangular shape) was significantly (97%) less cytotoxic than 3R4F smoke and that for deposited mass Vype ePen aerosol was significantly (94%) less cytotoxic than 3R4F smoke. Finally, based on the estimated deposited nicotine the Vype ePen aerosol was significantly (70%) less cytotoxic than 3R4F smoke. Thus, their work showed a reduced cytotoxic effect of Vype ePen aerosol ranging from 70% to 97%. In our study we observed that NCI-H292 cells exposed to 5 puffs of 1R6F undiluted smoke using the HCI regime had a significant reduction of viability (26.45% viable cells) whereas exposure of cells to undiluted Vype ePen3 aerosol under mHCI regime has a 97.57% of viable cells, with a reduced cytotoxic effect of >71%. Moreover, we evaluated the cytotoxic effect of the Vype eStick aerosol produced under the CRM81 regime (puff volume, duration and frequency of 55 ml, 3 s, 30 s, (55/3/30) and with a rectangular shape), obtaining a reduced cytotoxic effect of >74%. These evidences confirmed the results obtained by Azzopardi (Azzopardi et al., 2016) also by exposures to undiluted smoke and aerosol, supporting the reduced potential of e-cigs relative to conventional cigarettes in an *in vitro* model of bronchial epithelial cells even under more extreme exposure conditions (undiluted aerosol) than those of replicated papers.

The main results obtained by Azzopardi (2015, 2016) and Jaunky (2018) proved to be reproducible, thus brings more confidence that the conclusions on cytotoxicity derived from these results are true. Contrariwise, results on inflammatory and remodeling markers were non-reproduced. This does not automatically mean that conclusions derived from the result of original papers are false, but it does mean that conclusions and/or methods that were used in the two papers, including this, should be reconsidered and matter of further investigations. Surely, we believe that normalization of the raw data with viable cells would be appropriate for a more proper interpretation of data on inflammatory mediators, because they are actively produced and released by cells as a response to an exogenous stimulus. Despite this, our normalized data leaves us perplexed with respect to the data expected from the scientific literature (Strzelak et al., 2018; Yang et al., 2006). Undoubtedly, normalization of data with more accurate data on cell functionality, rather than viability, would provide more correct data. This aspect has to be taken into account for future evaluations in this direction.

In our opinion this replication study can allay doubts about results and proper execution of these three relevant studies confirming cytotoxicity of ENDS compared to cigarette smoke on bronchial epithelial cells improving scientific knowledges on this matter. Working with a multicentric approach the results here obtained has a high degree of agreement. An additional important milestone achieved with this study, was also the establishment of collaboration between the scientists of the Centers involved in this multi-center study that will allow us to create new teamwork aimed at the progression of knowledge in the field of reduced risk no-burning products.

## REPLICA PROJECT GROUP

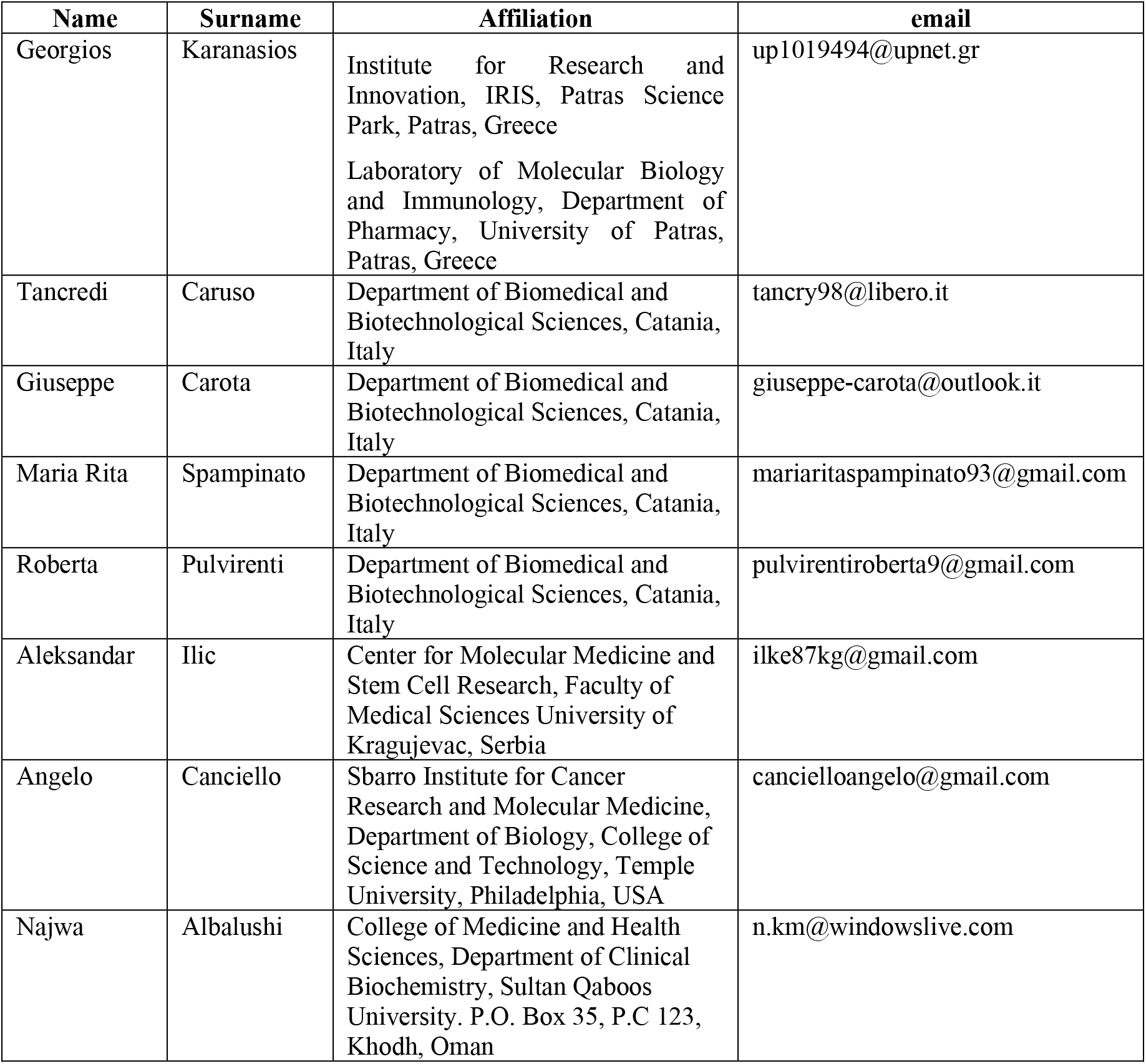

